# Student-led experimental evolution reveals novel biofilm regulators of adaptation to multiple niches

**DOI:** 10.1101/2025.06.06.658356

**Authors:** Abigail M. Matela, Colton W. Siatkowski, Changhua Yan, Sachin Thiagarajan, Vaughn S. Cooper

## Abstract

We established a research-education partnership, EvolvingSTEM, that currently provides thousands of secondary school students the opportunity to conduct authentic research experiments centered on microbial evolution each year. Providing high school students access to research experiences not only improves learning and can have positive and long-lasting impacts on their attitudes towards science, but also gives them the opportunity to make impactful scientific contributions. Through EvolvingSTEM, students evolve populations of *Pseudomonas fluorescens* in a bead model that includes daily cycles of bacterial dispersal, attachment, and biofilm growth and observe heritable changes in colony morphology. Genome sequencing of 70 mutants that they picked identified parallel mutations in genes known to regulate biofilm growth (*wsp*, *yfiBNR*, *morA, fuzY*) and uncovered novel adaptations: loss-of-function mutations in phosphodiesterase PFLU0185 that did not alter colony morphology and mutations affecting periplasmic disulfide bond formation producing small colonies. PFLU0185 mutants rapidly and consistently reached high frequencies and phenotyping revealed roles in cyclic di-GMP regulation, biofilm formation, and motility, prompting us to name this gene *bmo* (<u>b</u>iofilm and <u>mo</u>tility <u>regulator</u>). Competition experiments and microscopy demonstrated *bmo* mutants employ generalist strategies and coexist with their ancestor and specialist mutants through niche differentiation. Consequently, phenotypic diversity is maintained, with smooth (ancestral and *bmo*) colonies consistently outnumbering wrinkly and fuzzy variants. This study advances our understanding of biofilm genetic architecture while demonstrating that student-led research can uncover mechanisms of microbial adaptation relevant to *Pseudomonas* infection biology.

**IMPORTANCE:** Bacterial biofilms dominate microbial life, yet their evolutionary genetics remain incompletely understood. Extensive replication of experiments that employ similar, but not identical, biofilm selection models can provide valuable insights into mechanisms of adaptation. We demonstrate that this can be achieved through university-education partnerships that engage secondary school students in authentic research. Student-led experiments revealed that loss-of-function mutations in a conserved phosphodiesterase, PFLU0185/*bmo*, dominate evolved populations without changing colony morphology. This finding, combined with diverse, less frequent mutants that alter colony morphology informs the process of biofilm niche differentiation. This work also demonstrates the power of distributed research networks for discovering new genetic pathways of adaptation. Students gained authentic research experience, potentially inspiring them to become scientists, while identifying mutants adapted to discrete conditions that maintain diversity within biofilms. This synergy between education and discovery offers a scalable model for addressing complex biological questions while developing scientific literacy in diverse classrooms.

## INTRODUCTION

Despite being the dominant form of microbial life, our understanding of the ecology, evolution, and genetic controls of adaptation within biofilms is still developing. What we and many others have learned is that bacteria evolve rapidly and conspicuously, often by producing new colony morphologies, when selected to improve surface growth in experimental models as well as in clinical and environmental settings (Stewart & Franklin, 2008; Poltak & Cooper, 2011; Traverse et al., 2013; Ellis et al., 2015; Nadell et al., 2016). This dynamic can be captured in simple experiments conducted by secondary school students that not only improves their learning experiences but also provides them the opportunity to contribute to cutting-edge scientific research (Cooper et al., 2019; Spiers, 2014; Taylor et al., 2024).

We might expect that interactions in biofilms would be complex as cells engineer their environment by secreting polymers that bind them to a surface and protect them from stresses, competitors, or predators. Two overlapping processes can explain the origin and maintenance of phenotypic diversity within biofilms. The first is environmental structure itself, which allows multiple phenotypically equivalent mutants to arise and coexist in different regions (Kassen & Rainey, 2004). The second is ecological interactions made possible by structure, for example, when an early adhering population facilitates the attachment of different types (Habets et al., 2006; Harris et al., 2021). However, the genetic and phenotypic outcomes of these processes are less clear but broadly important in natural and engineered systems, such as wastewater treatment (Røder et al., 2016) and in the clinic (Trubenová et al., 2022). For example, high-biofilm forming bacterial and fungal mutants often arise during chronic infections, are associated with increased resistance to antibiotics and immune phagocytes, and worsen patient outcomes (Hall-Stoodley et al., 2004; Flemming & Wuertz, 2019; Pestrak et al., 2018). There is a need to broadly characterize the spectrum of genotypes and phenotypes that generate biofilm diversity. Experimental evolution of microbial populations under conditions favoring biofilm combined with contemporary genomic analysis can be a powerful screen of these traits.

Our laboratory previously developed an *in vitro* model to study evolution throughout the entire biofilm life cycle of surface attachment, biofilm assembly, dispersal, and re-colonization (Poltak & Cooper, 2011). In short, bacteria are cultured in test tubes that contain growth media and a polystyrene bead, which serves as a surface for bacteria to colonize and form a biofilm. The biofilm-covered bead is then transferred to a new test tube with fresh media and a sterile bead. Beads are serially transferred to select for biofilm-adapted mutants that can disperse from the old bead and recolonize and assemble a new biofilm on the new bead every 24 hours. This model has been used by us to better understand the dynamics of biofilm evolution and associated ecological diversification in multiple species (*Burkholderia cenocepacia*, *Acinetobacter baumannii*, *Pseudomonas aeruginosa*) under various environmental stressors (e.g., nutrients, antibiotics) (Poltak & Cooper, 2011; Traverse et al., 2013; Cooper et al., 2014; Ellis et al., 2015; O’Rourke et al., 2015; Flynn et al., 2016; Turner et al., 2018; Santos-Lopez et al., 2019; Scribner et al., 2020; Mhatre et al., 2020; Harris et al., 2021). The bead model has also been adapted by many others to address similar questions and is routinely paired with whole genome sequencing of mutant clones and evolved populations to identify causes of new phenotypes (Henriksen et al., 2022; Oakley et al., 2021; Trampari et al., 2021). A surprising finding from these studies is the extent of genetic parallelism rather than a diverse array of pathways to adaptation. Despite the intuition that the biofilm life cycle is both structured and heterogeneous, and the observation of various colony phenotypes, mutations in relatively few genes encoding major biofilm regulators accounted for most of the adaptations (Mhatre et al., 2020; Santos-Lopez et al., 2019).

One possible explanation for convergent evolution in the biofilm bead model is that each study was conducted by one or few investigators using well controlled conditions, thereby focusing selection on limited traits. Higher experimental replication by many different researchers could address this concern. For this reason and many others related to improving learning and access to authentic research, we adapted the bead model for use in secondary school classrooms (Cooper et al., 2019). In this research-education partnership, students work with the harmless plant probiotic bacterium, *Pseudomonas fluorescens* strain SBW25, which is both a model of this environmentally relevant species and shares many orthologs with the opportunistic pathogen *P. aeruginosa*. This program is called EvolvingSTEM and now engages thousands of students a year in authentic microbiology research in grades 6-12 in more than 20 schools in several U.S. states. Previous research on *P. fluorescens* SBW25 identified biofilm-related adaptations to static culture conditions, where populations rapidly evolve to form a biofilm mat at the oxygen-rich, air-liquid interface (Koza et al., 2017). The evolved populations diversified into three distinct colony morphologies: (1) smooth and round (the ancestral phenotype), (2) wrinkly, and (3) fuzzy (Rainey & Travisano, 1998). It is this rapid evolution to produce pellicles that encouraged us to use *P. fluorescens* with our bead model to study its adaptation to biofilm growth with students.

The most conspicuous new colony morphology seen in previous experiments is the wrinkly phenotype. This phenotype commonly results from mutations in one of three genetic pathways – *wsp*, *yfiBNR* (or *aws*), and *morA* (or *mws*) – that regulate the production of a secondary messenger, bis-(3’-5’)-cyclic dimeric guanosine monophosphate (cyclic di-GMP) (McDonald et al., 2009). Cyclic di-GMP controls biofilm formation by regulating the activity of several downstream genes and proteins (Jenal et al., 2017). The production of cyclic di-GMP is regulated by the opposing activity of two enzyme families: diguanylate cyclases (DGCs) that synthesize cyclic di-GMP and phosphodiesterases (PDEs) that degrade it (Jenal et al., 2017). High cellular levels of cyclic di-GMP are associated with surface attachment and biofilm production, while low levels are associated with increased motility, solitary behavior, and bacterial dispersal (Laventie, Benoit-Joseph, 2020). Mutations in the *wsp*, *yfiBNR*, and *morA* pathways lead to constitutive activation of a DGC, which results in overproduction of cyclic di-GMP (Malone et al., 2007; McDonald et al., 2009), and, ultimately, increased biofilm formation through increased production of acetylated cellulose (Spiers et al., 2002, 2003). In contrast, fuzzy phenotypes are a result of mutations in *fuzY*, which encodes a β-glycosyltransferase predicted to modify LPS O-antigens. Mutations in *fuzY* are thought to promote cell-cell contact but the mechanistic benefit of mutations in this gene are less well understood (Ferguson et al., 2013).

*P. fluorescens* populations evolved by high school students with our EvolvingSTEM evolution-in-action experiment often exhibit diverse colony morphologies within and among populations that resemble those seen under selection in static culture conditions (Cooper et al., 2019). We wondered if the phenotypic similarities of mutants adapted to static culture and those in our bead model are caused by the same underlying genetic changes, so we sequenced 70 clones from populations almost entirely evolved by high school students. We found that *P. fluorescens* adapts to form biofilms on surfaces through mutations in some of the same genetic pathways as those used to form pellicles in unshaken vials and also uncovered novel biofilm adaptations. We then sought to identify what drives evolutionary dynamics in our bead model and whether these dynamics differ when bacteria are grown in subinhibitory levels of the common antimicrobial triclosan, which was used to prevent fungal and bacterial contamination at some schools. This led to the discovery that biofilm adaptation in our model is primarily driven by lineages that do not change colony appearance but acquire loss-of-function mutations in PFLU0185, a highly conserved gene across pseudomonads encoding a DGC and dominant PDE (Cai et al., 2022, 2022; Eilers et al., 2022; Eilers, Yam, et al., 2024; Wei et al., 2019). Although PFLU0185 mutants share the same colony morphology as their ancestor, further phenotyping revealed adaptations in biofilm formation and motility. We therefore suggest renaming this locus *bmo*, for <u>b</u>iofilm and <u>mo</u>tility regulator. Population sequencing, microscopy, and multi-genotype competition experiments suggest that *bmo* mutants coexisted with their ancestor and with rarer, genetically diverse lineages producing wrinkly and fuzzy colonies in dynamics that can be explained by multi-niche selection (Brisson, 2018).

## RESULTS

### Student-led experiments rapidly uncover diverse phenotypes

Our EvolvingSTEM evolution-in-action experiment allows students to observe bacterial populations evolving and diversifying within days as they adapt to a daily selection regime to complete the entire biofilm life cycle of dispersal, surface attachment, and biofilm maturation. Students work with a harmless plant probiotic bacterium, *P. fluorescens* strain SBW25. As the populations evolve, students can observe at least two types of phenotypic change. First, as high biofilm-forming mutants increase in frequency in the population, students often see a more robust layer of biofilm form in their test tubes at the air-liquid interface. Second, students can observe changes in the colony morphologies of bacteria grown on agar plates. While the ancestral population produces only smooth, circular colonies, evolved populations often exhibit diverse, conspicuous colony morphologies within and among populations (Fig. 1).

**Fig. 1.**
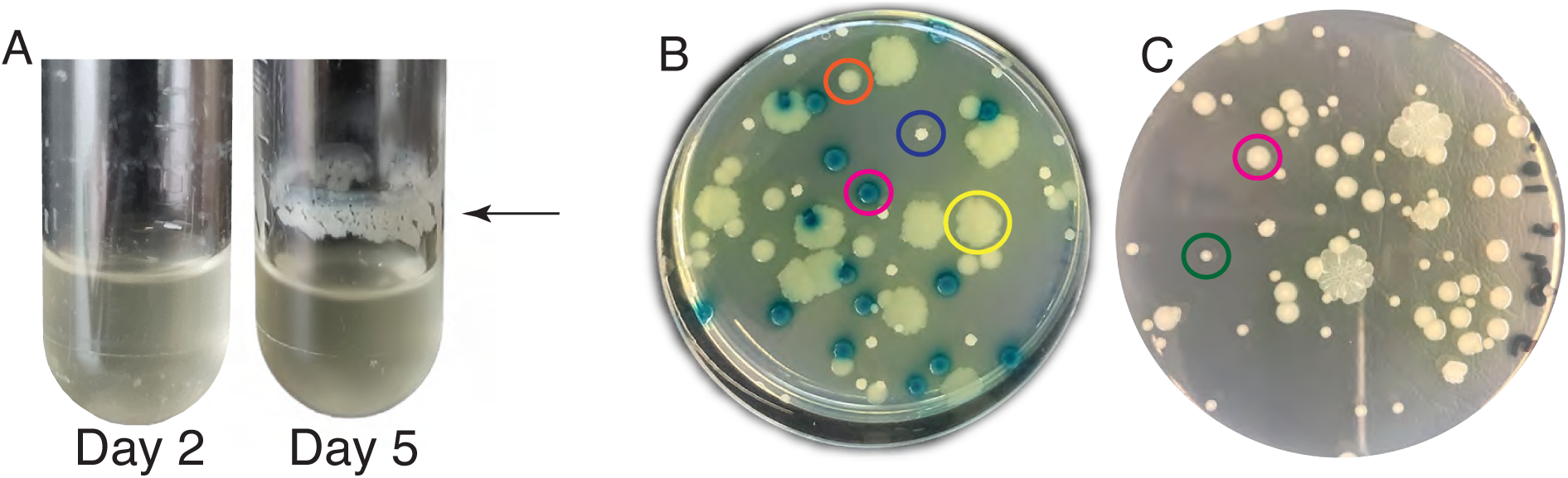
Biofilm growth patterns and colony morphologies of *P. fluorescens* SBW25 and evolved biofilm-adapted mutants. Evolved populations often have a distinct ring of biofilm at the air-liquid interface (A). The *P. fluorescens* ancestor (lac+) has blue, smooth-edged circular colonies (pink circle), the *bmo* (PFLU0185) mutant has yellow, smooth-edged circular colonies (orange circle), the *fuzY* mutant has large, yellow, irregular colonies (yellow circle), and the *wspF* mutant has small, white, irregular colonies (purple circle) (B). Small colony variants (green circle) can be distinguished from their ancestor (pink circle) by their significantly smaller size (B).

Students from three high schools – Winnacunnet High School (WHS, Hampton, NH, N=∼250), Pittsburgh Science and Technology Academy (Pittsburgh, PA, N=25), and Allderdice High School (Pittsburgh, PA, N=24) – conducted experiments under slightly different conditions. Changes to the experimental protocol were made to accommodate class requirements and increase the accessibility of EvolvingSTEM in varied classroom settings (our updated protocol can be found in supplemental materials). While WHS used standard aseptic technique, in other schools, we added the commonly used antimicrobial compound triclosan to the media and agar plates to broadly inhibit bacterial and fungal contamination. *Pseudomonas* species are known to tolerate or resist triclosan (Schweizer, 2001; McFarland et al., 2021), and this eliminates the need to use a flame in schools that do not have access to Bunsen burners and/or do not feel comfortable having students work near an open flame. Another difference was that WHS aerated growing bacterial cultures on a roller drum, which was not available in most classrooms, so we switched to using orbital shakers to aerate growing cultures.

Students plated samples of their bead-attached bacterial populations on agar at the beginning and end of the experiment. Ancestral *P. fluorescens* clones produce only smooth, round colonies, and this was the only morphology identified on the early plating day. Most bead-adapted populations contained wrinkly colony variants on the final plating day, and some had fuzzy and small colony variants (Fig. 1). Wrinkly and fuzzy colony variants had a similar appearance to those identified when *P. fluorescens* is under selection for the formation of biofilm mats in static culture conditions (Koza et al., 2017). While our experimental design also selects for biofilm-adapted mutants, it differs from static selection experiments in that bacteria must colonize a new surface each day, completing the entire biofilm life cycle of dispersal, attachment, biofilm assembly, and matrix building. We therefore explored if the convergent colony phenotypes we identified in these experiments was the result of convergent mutational targets by sequencing the genomes of wrinkly and fuzzy clones from independent populations. In addition, we sequenced the genomes of smooth colonies, which were identical in appearance to the ancestor, and small colony variants, which were the only other unique morphology noted in these experiments (Fig. 1).

### Biofilm adaptations include mutations affecting cyclic di-GMP production, LPS O-antigen structure, and periplasmic protein folding

Two remarkable properties of this collection of mutant genomes deserve highlighting before we explore the genes affected or their functional implications. First, in 55 of 70 mutants, only one mutation was identified, meaning that this single mutation experienced strong selection to rise to a detectable frequency and, in most cases, change the colony morphology (Table 1, Table S1). Consequently, the biofilm bead model serves as a potent forward genetic screen for single mutations that influence biofilm-related fitness. Of the remaining 15 mutants, 13 had two mutations and one had three mutations. Second, the extent of site-specific parallelism in these mutants is extraordinary: 13 individual mutations occurred more than once in different clones. Although it is possible that a few instances of parallel mutations were not entirely independent, e.g. from the same school and year, most clearly arose independently from different schools and years. Exceptional examples include one SNP in *wspF* that was observed in six clones and another 33bp deletion in *yfiR* observed five separate times (Table 1). This observation suggests extremely strong selection for certain mutations, a locally higher mutation rate, or both. Overall, 23 unique genes became mutated and were affected by a roughly equal mix of small insertion/deletions (indels) and SNPs with fewer larger deletions. This spectrum of mutations demonstrates that *P. fluorescens* experiences strong selection for a narrow set of traits in the biofilm bead model that often favors identical mutations in independent experiments.

**Table 1.**
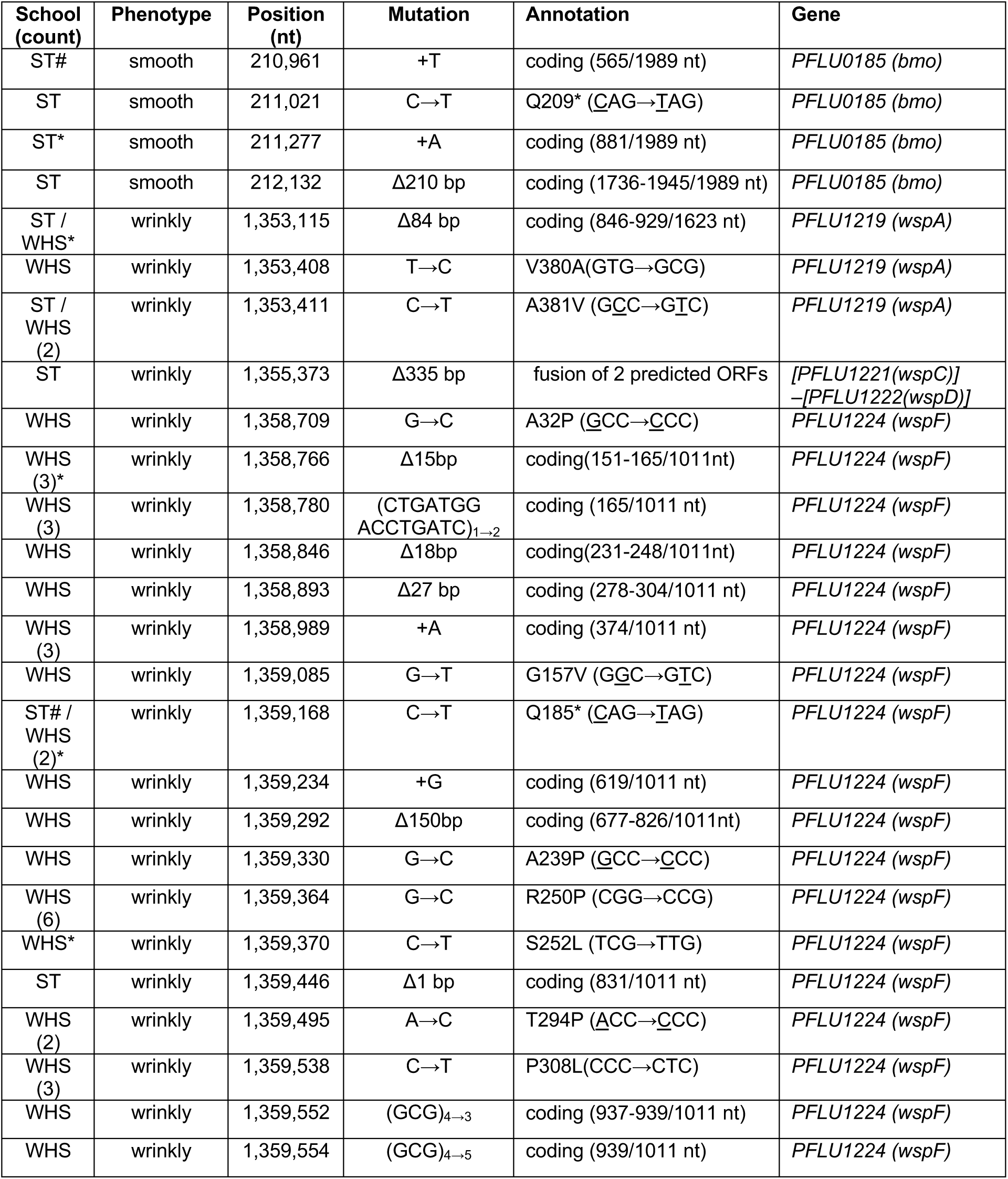
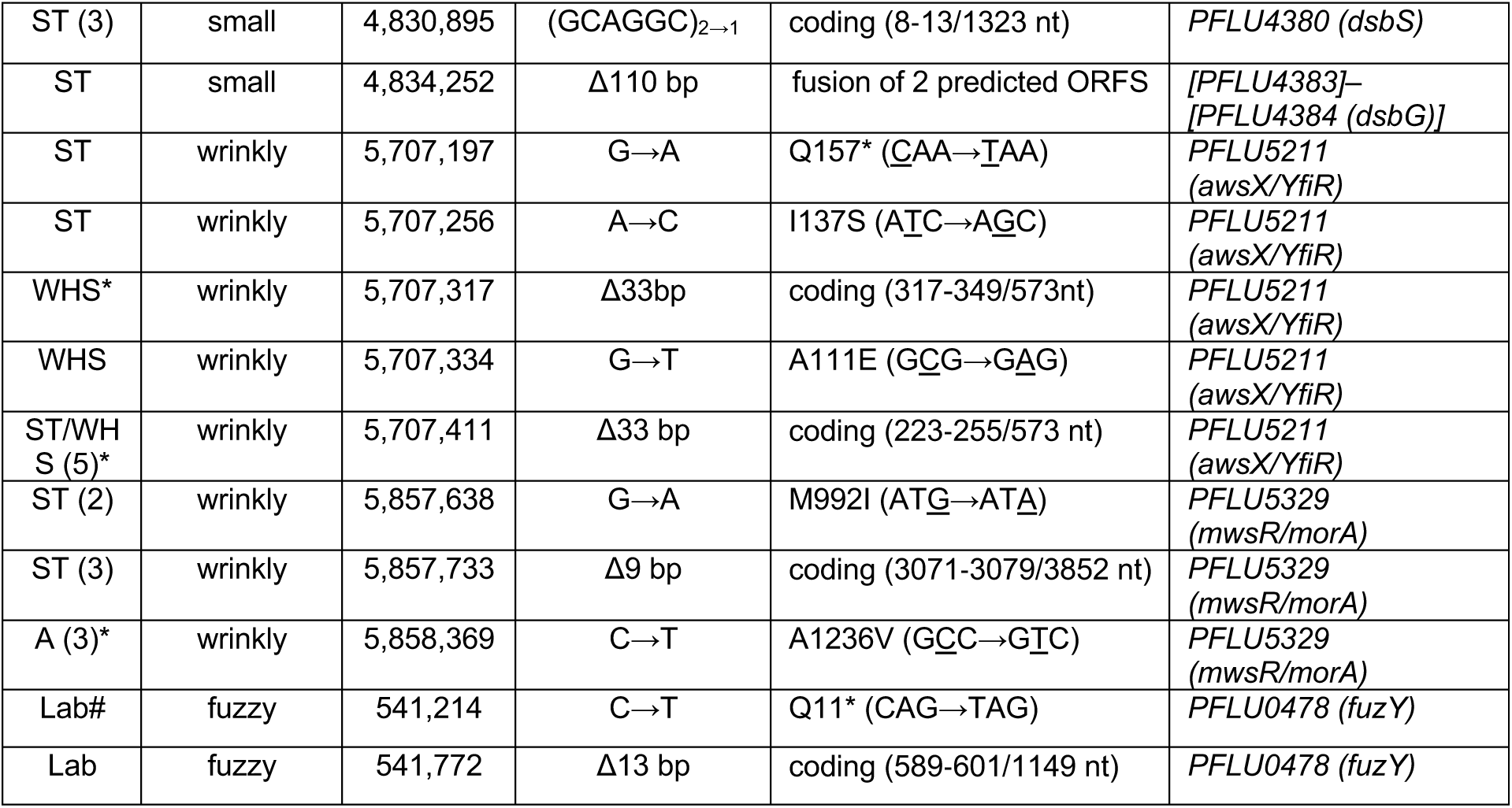
Genotypes of *P. fluorescens* mutants determined by WGS that were isolated based on colony phenotype by students in different schools (ST, WHS, A, or Lab = our research lab). *denotes presence of secondary mutations, see Table S2 for details. #denotes representative mutant used in phenotypic studies

The genomes of 57 clones with wrinkly phenotypes were sequenced from 50 independently evolved biofilm populations. In agreement with previous studies on biofilm formation in static culture conditions, causative mutations were identified in the *wsp*, *yfiBNR*, and *morA* pathways (Fig. 2, Table 1), which supports a critical role for these pathways in biofilm regulation under diverse selective regimes. Mutations in the *wsp* pathway were the most common cause of wrinkly phenotypes across all sequenced clones, with approximately 70% having a single causative mutation in this pathway. Mutations were primarily found in *wspF* (PFLU1224, 33 clones), followed by *wspA* (PFLU1219, 6 clones). In addition, a single clone had a 335 base-pair deletion that overlapped the coding regions of *wspC* (PFLU1221) and *wspD* (PFLU1222), resulting in a WspC-WspD fusion protein. The *yfiBNR* pathway was the second most common cause of wrinkly phenotypes, with approximately 16% of clones having a single causative mutation in *yfiR* (PFLU5211, 9 clones). Mutations in *morA* (PFLU5329, 5 clones) were the least common cause of wrinkly phenotypes, with approximately 9% of clones having a single causative mutation. Surprisingly, all three clones isolated at Allderdice had mutations in two pathways. These clones contained a single, nonsynonymous SNP in the PDE domain of *morA*. One clone had an additional nonsense mutation in *wspF*; one had an additional missense mutation in *wspA*; and one had an additional deletion in the linker region between the DGC and PDE domains of *morA*. These mutants indicate that the sequential acquisition of multiple mutations in pathways influencing cyclic-di-GMP metabolism can be beneficial.

**Fig. 2.**
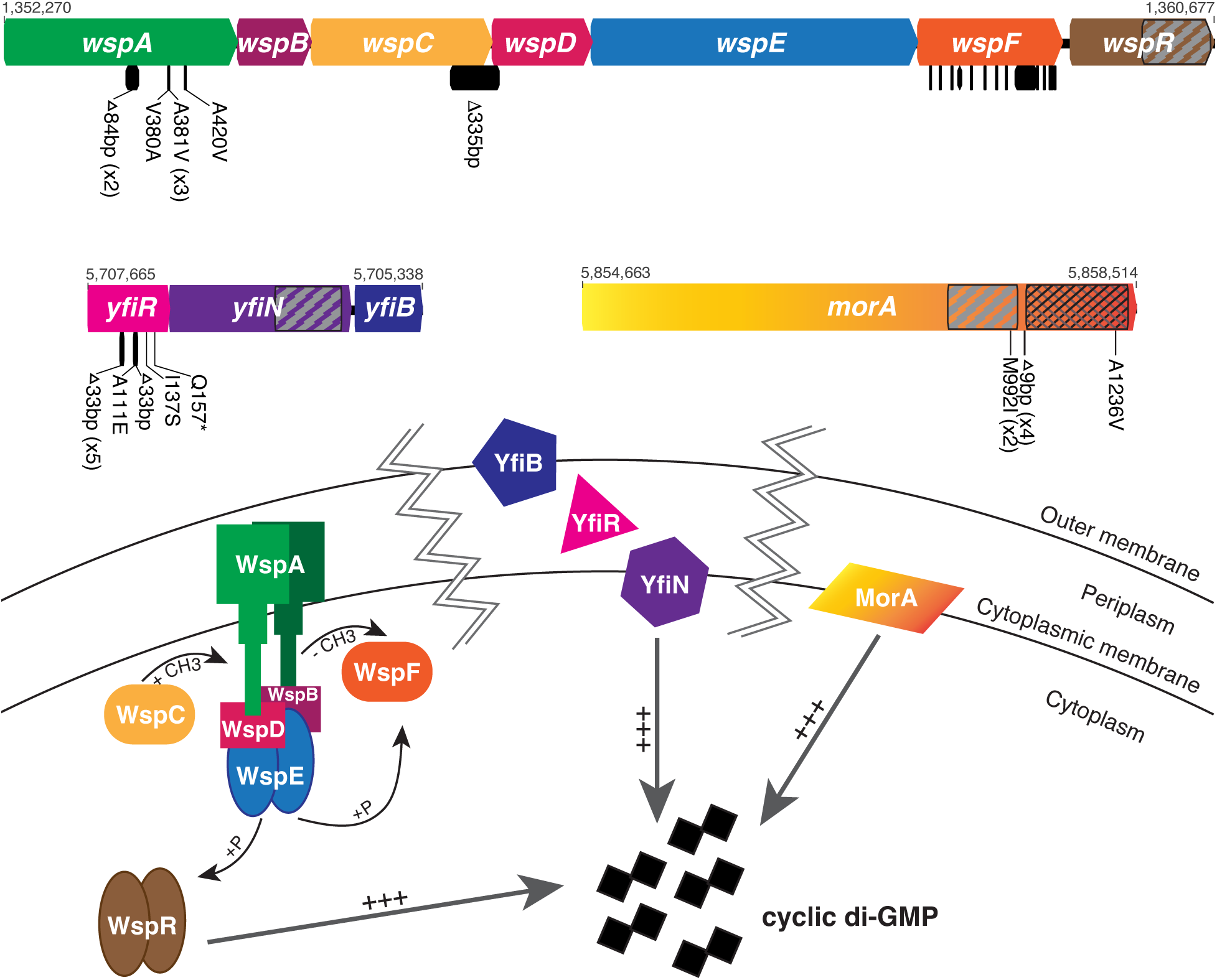
Wrinkly colony phenotypes are caused by mutations in the *wsp*, *yfiBNR*, and *morA* pathways. Mutations are shown as black bars below the gene diagrams. When multiple independent mutations occurred, their counts are in parentheses. Mutations in *wspF* (31 in total) are too numerous to display. DGC domains are indicated with diagonal hatching; PDE domains are indicated with cross hatching. Genome positions of the beginning and end of each operon/gene are indicated above the gene diagrams. Table 1 and Table S1 contain a complete list of mutations. All mutations are predicted to cause constitutive activation of a DGC, increasing cyclic di-GMP synthesis.

Fuzzy mutants were identified in evolved populations at several schools, although less frequently than wrinkly mutants. While we did not sequence any fuzzy mutants from high school experiments, two fuzzy mutants isolated in the laboratory had predicted loss-of-function mutations in *fuzY* (PFLU0478) (Fig. 3). In addition, we sequenced the genomes of five small colony variants isolated by students and in the laboratory. These mutants contained single mutations in a predicted two-component system sensor histidine kinase, *dsbS* (PFLU4380), or in the operon it is predicted to regulate, *dsbD* (PFLU4382)-*dsbG* (PFLU4384) (Fig. 3). Three clones had an in-frame deletion in the N-terminal region of *dsbS*, one clone had a 110 base-pair deletion that overlapped the coding regions of *dsbE* (PFLU4383) and *dsbG*, resulting in a DsbE-DsbG fusion protein, and one clone had a mutation in the intergenic region upstream of *dsbR* (PFLU4381) and *dsbD* (Fig. 3). Lastly, we also sequenced clones from five evolved populations that retained the ancestral smooth morphology. Although one smooth clone had no mutations, four clones had a mutation in a gene encoding both DGC and PDE domains, PFLU0185, and this was the only mutation identified in three clones (Fig. 3). We have named this gene *bmo* for its roles in biofilm formation and motility, which are further described below.

**Fig. 3.**
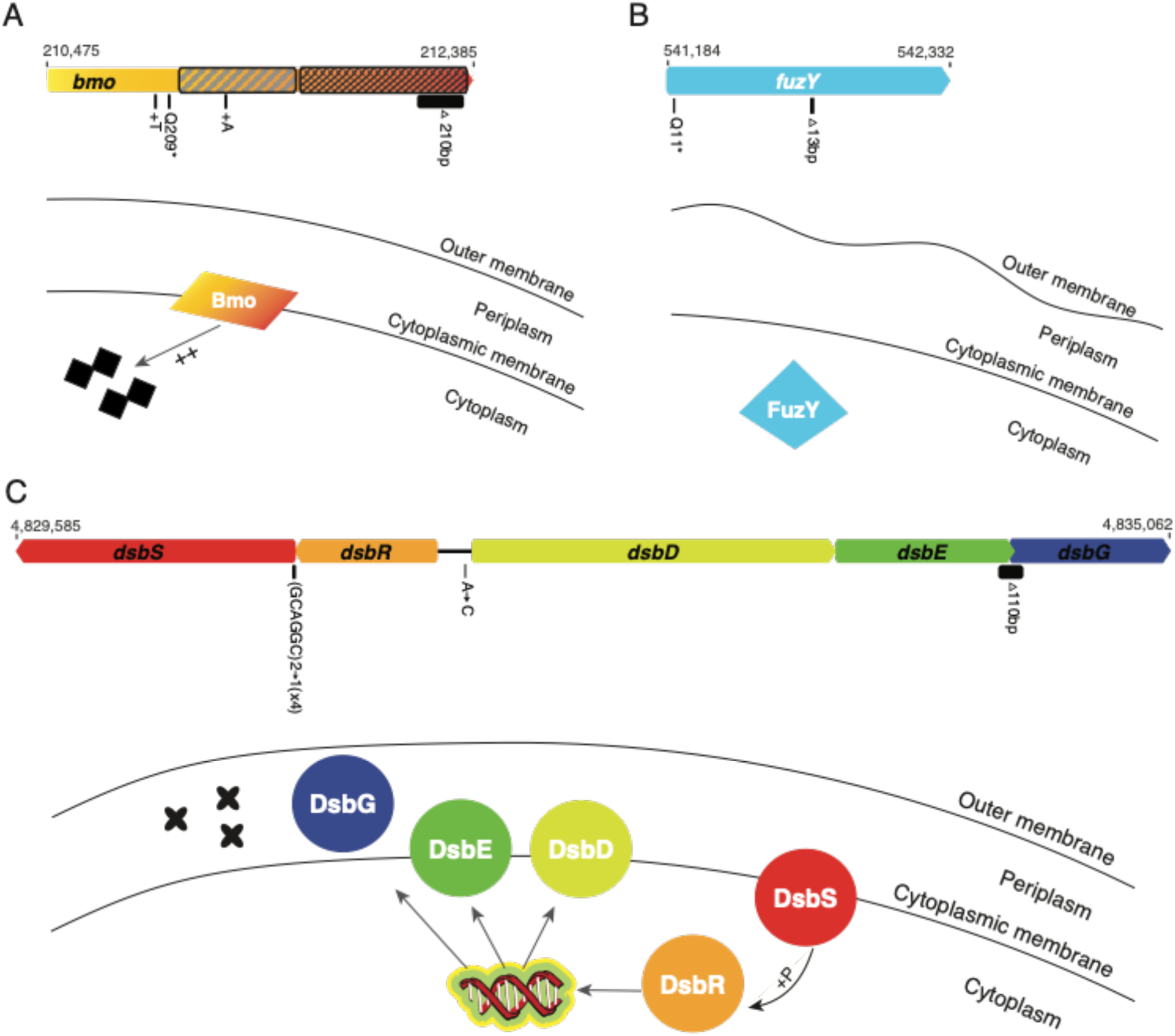
Biofilm adaptations caused by mutations in LPS, disulfide bond formation, and a novel regulator of cyclic-di-GMP. Mutations are shown as black bars below the gene diagrams. When multiple independent mutations occurred, their counts are in parentheses. DGC domains are indicated with diagonal hatching; PDE domains are indicated with cross hatching. Genome positions of the beginning and end of each operon/gene are indicated above the gene diagrams. Mutations in *bmo* (PFLU0185) are predicted to increase basal levels of cyclic di-GMP (A). Mutations in *fuzY* are predicted to modify LPS O-antigens (B). Mutations in *dsb* genes are predicted to cause misfolded periplasmic proteins. Table 1 and Table S1 provide mutation details.

### Strong selection for mutations in *bmo* during adaptation to the experimental biofilm life cycle

We were intrigued by the discovery of mutations in a DGC-PDE encoding gene that did not change the mutant colony phenotype. Although an objective of the EvolvingSTEM program is to visually demonstrate adaptation by natural selection by observing novel colony phenotypes after selection for improved biofilm formation, in fact, most populations have a majority of colonies that retain the smooth appearance of the ancestor after one week of passage. This suggests that focusing only on colony variants may miss biofilm adaptations. To address this limitation, we conducted a 15-day evolution experiment in which we used whole-population, whole-genome sequencing to identify all mutations reaching appreciable frequency in biofilm populations regardless of whether they alter colony appearance. We also replicated this comparison in media containing triclosan, which we use in classrooms to guard against contamination and which appears to accelerate rates of diversification. Specifically, four replicate biofilm populations were propagated with and without 25µg/mL triclosan for 15 days. We plated the bead-attached portion of each population after 4, 8, 11, and 14 transfers to observe phenotypic changes to population colony morphologies over time and performed whole-population, whole-genome sequencing after 2 and 14 transfers (Table S2, Fig. 4). We noted three distinct colony morphologies, wrinkly, fuzzy, and small, in addition to the ancestral, smooth phenotype. Populations grown in triclosan had moderately increased diversity on the earliest plating day, but this diversity decreased over the course of the experiment, with all populations having over 95% smooth colonies by the final plating. Populations that were grown without triclosan maintained slightly higher diversity over time, and all populations had between 90-95% smooth colonies by the final plating (Fig. 4).

**Fig. 4.**
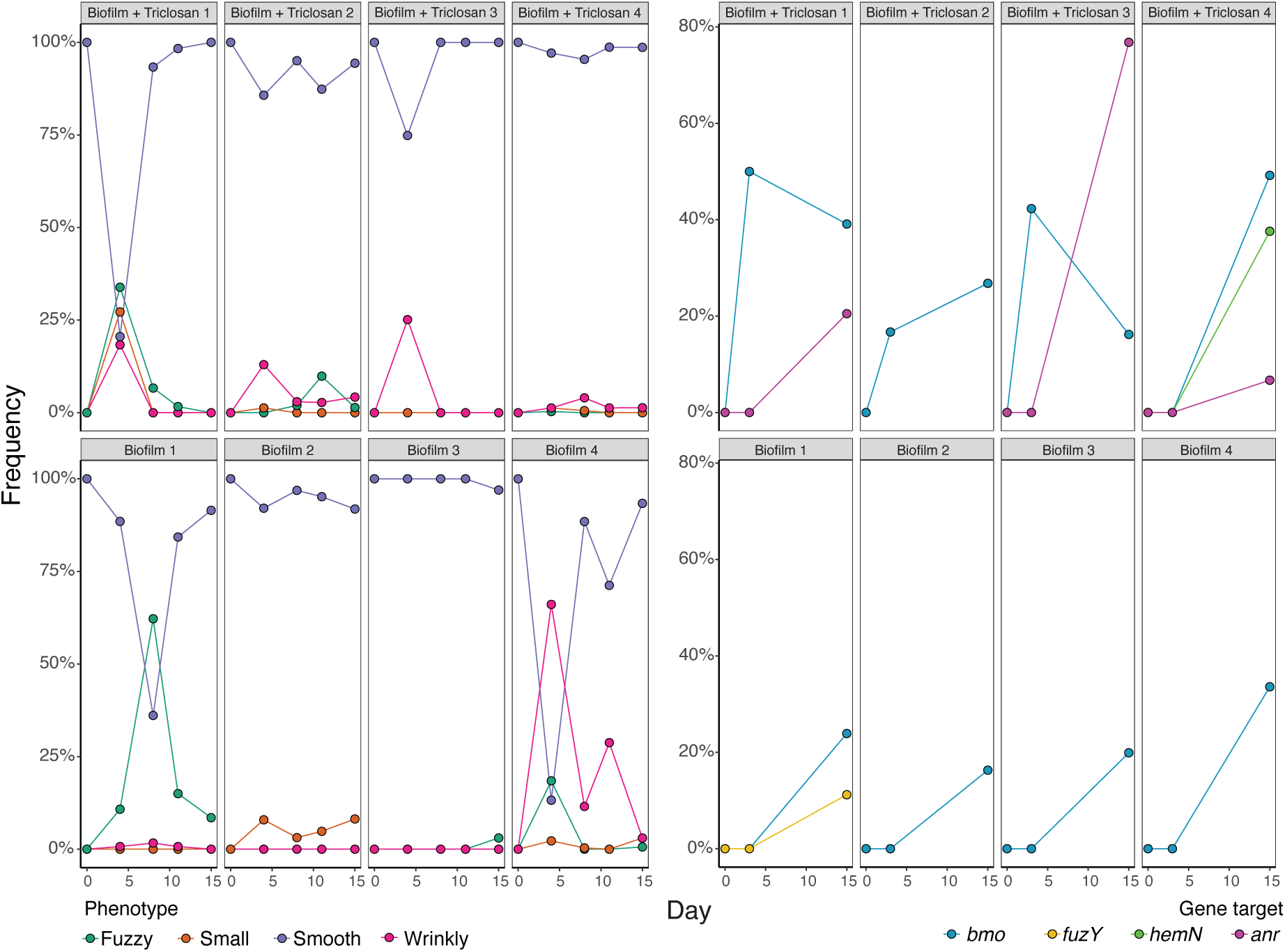
Dynamics of evolved phenotypes and genotypes during 15 days of experimental biofilm selection. Populations were grown for 15 days in QB media with or without triclosan with four plastic beads and passaged every 24 hours by transferring one bead into fresh media with three new beads. Bead-attached populations were plated on days 4, 8, 11, and 15 and contained colonies with fuzzy, small, smooth, and wrinkly phenotypes (A). Whole-genome, whole-population sequencing was performed on days 3 and 15. Points include the cumulative frequency of mutations identified in each gene (B). A complete list of mutations can be found in Supplemental Table 1.

Surprisingly, mutations in the *bmo* locus were the only gene-level parallel genetic change in all biofilm populations, and they rose to detectable frequencies by the third day of the experiment in media containing triclosan, but not in media without triclosan (Fig. 4, Table S2). Multiple independent mutations in *bmo* were present within each of the three biofilm populations with added triclosan at day 3 at 16.7%, 42.3%, and 50% cumulative frequency, and by day 15 all eight biofilm-adapted populations had *bmo* mutants at >16% cumulative frequency. The only other notable biofilm-related mutation that we observed was in *fuzY*, which reached a frequency of 11.2% in one population that was grown without triclosan. Furthermore, three of the four populations grown with added triclosan gained mutations in a transcription factor, *anr* (PFLU4570), and one of the genes it regulates, *hemN* (PFLU4568). These mutations may be adaptations to growth in triclosan and will be the subject of future study.

### Mutants are differentiated in biofilm phenotypes including attachment, assembly, and motility

We predicted that mutations in *bmo* achieved the highest frequencies and repeatedly evolved in our biofilm selection model because they are best adapted to the cyclical requirements to attach to the plastic bead, assemble biofilms, and then disperse to colonize a new bead. We therefore characterized the motility, biofilm production, and cyclic di-GMP production for a representative *bmo* loss-of-function mutant with a single nucleotide insertion that results in a premature stop codon truncating both the DGC and PDE domains (Table 1). We compared its biofilm associated phenotypes to a representative wrinkly (*wspF* Q185*) and fuzzy (*fuzY* Q11*) mutant, along with the *P. fluorescens* SBW25 ancestor from which these mutants evolved.

We first compared three common types of bacterial motility – swarming, swimming, and twitching. All biofilm-adapted mutants had reduced swarming and swimming motility in comparison to the ancestor (Fig. 5A, 5B). *wspF* mutants were the least motile, while *fuzY* and *bmo* mutants had intermediate phenotypes. Twitching motility was not displayed by any biofilm-adapted mutants or the ancestor, confirmed by comparison to *Pseudomonas aeruginosa* PAO1 which possesses twitching motility (Fig. S1).

**Fig. 5.**
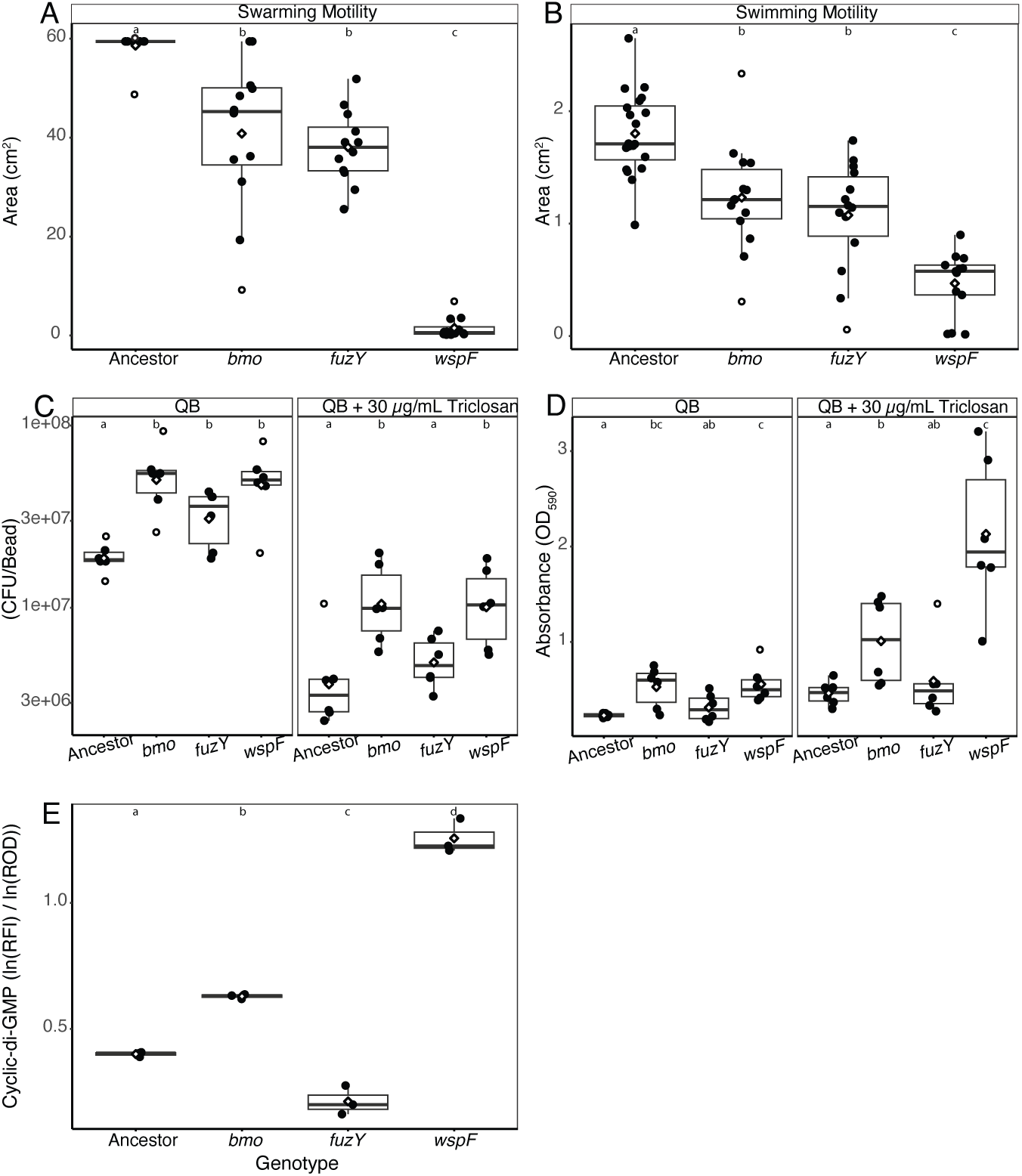
Biofilm-related phenotypes of *P. fluorescens* mutants. Swarming motility (A) and swimming motility (B) show reduced levels compared to the ancestor. Biofilm production measured by attached colony forming units (C) and by crystal violet absorbance at 590nm (D). Points are the average of three measurements recorded for each replicate. Results in QB media are similar but more pronounced with the addition of triclosan. Relative cyclic di-GMP production from a fluorescent reporter (E) is reported as ln(RFI)/ln(ROD600), where RFI is relative fluorescence intensity and ROD600 is relative optical density. In all panels, mean values are indicated by diamonds, and outliers are indicated by an open circle. Statistical comparisons were conducted by one-way ANOVA with a Games-Howell post hoc test with p = 0.05, with letters at the top of each panel indicating statistically distinct groupings.

To estimate biofilm production by each mutant, we adapted the traditional crystal violet 96-well plate assay (O’Toole & Kolter, 1998) to better align with how bacteria form biofilms in our bead model. Bacteria were cultured for 24 hours in test tubes containing media and two plastic beads. We quantified total biofilm production by staining one bead with crystal violet, removing excess stain by washing the bead in water, and solubilizing biofilm-bound crystal violet by vortexing the bead in solvent. We quantified biofilm population size by vortexing and dilution-plating from the second bead. When grown in QB medium, *bmo* and *wspF* mutants have the highest total biofilm production and population size (Fig. 5C, 5D). One caveat to note is that *wspF* mutant populations often acquire compensatory, suppressor mutations that can rise to high frequencies in overnight cultures, so between 20%-40% of the colonies counted had a smooth, ancestral phenotype. When grown in media with added triclosan, total biofilm production is doubled in *bmo*, *fuzY*, and the ancestor and quadrupled in *wspF* (Fig. 5C, 5D). Moreover, the population size of all genotypes is less than half that observed in triclosan-free media (Fig. 5C, 5D). Planktonic growth is also slower when triclosan is included in the media and final population sizes are smaller (Fig. S2).

To determine how these biofilm-adapted mutations affect cyclic di-GMP metabolism, we used a GFP reporter plasmid that responds to the intracellular concentration of cyclic di-GMP to estimate these changes. Levels of cyclic di-GMP differed significantly among *P. fluorescens* mutants following 24 hours of growth in QB media (Fig. 5E). Cyclic di-GMP concentrations were highest for *bmo* and *wspF* mutants, with *wspF* producing nearly double the amount of cyclic di-GMP as *bmo*. Surprisingly, the *fuzY* mutant produced approximately half as much cyclic di-GMP as the ancestor, although this gene is not reported to have a role in the regulation of cyclic di-GMP. Taken together, these results demonstrate that *bmo* regulates biofilm formation and motility and loss-of-function mutations in this gene allow for the optimization of biofilm production and planktonic growth in our bead model.

We sought to understand how the different macroscopic and regulatory phenotypes of *P. fluorescens* mutants generated distinct microscopic biofilm phenotypes. We therefore used confocal microscopy to image biofilms produced by our representative *bmo*, *wspF*, and *fuzY* mutants as well as the ancestor that contained a constitutively active fluorescent reporter inserted at a neutral site on the chromosome. Both the ancestor and *bmo* formed relatively confluent biofilms of even thickness and their similar appearances suggest that they would compete for the same niche space on the bead (Fig. 6). In contrast, the *fuzY* biofilm was more clumped and uneven in thickness and coverage, whereas the *wspF* biofilm was surprisingly sparse, with small, dense clusters (Fig. 6). It is notable that we imaged biofilms on glass slides rather than on the polystyrene beads on which these mutants evolved, and that the supernatant of the *wspF* cultures contained large aggregates that did not adhere well to glass. To explore how these mutants interact in a diverse biofilm, we imaged mutants with different fluorescent markers that were co-cocultured in pairs (Fig. S3). Overall, the observed patterns of biofilm assembly agreed with predictions from mutant growth alone, with interspersed and confluent growth by *bmo* and the ancestor but fewer discrete patches of different sizes and forms produced by *wspF* and *fuzY.* Taken together, these images indicate that biofilm mutants in different pathways are ecologically distinct and produce clusters or coatings of varying biomass and form, both alone and in mixture.

**Fig. 6.**
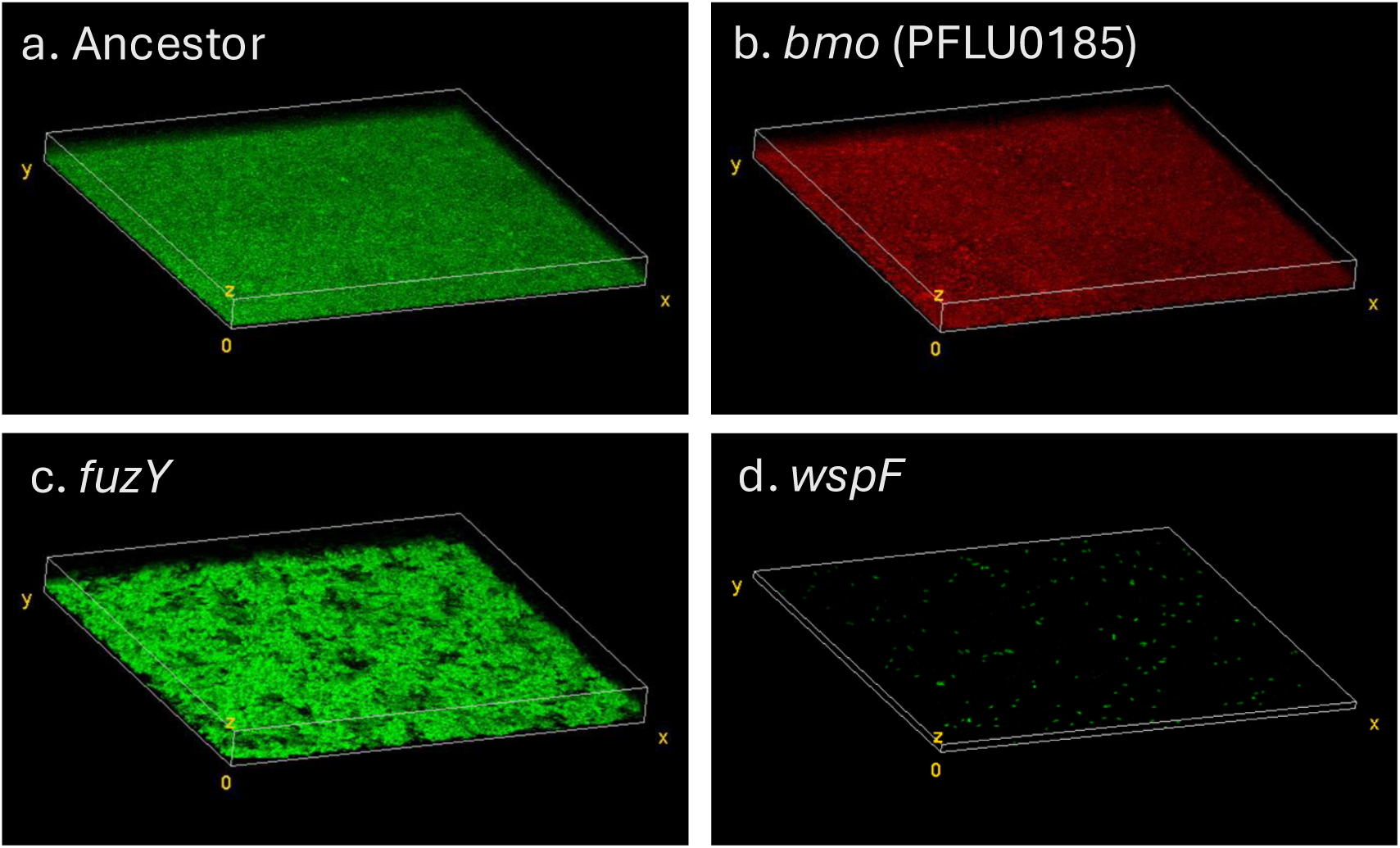
Confocal microscopy of representative *P. fluorescens* mutants shows differentiated biofilm phenotypes.. Ancestor (a), *fuzY* (c) and *wspF* (d) are tagged with eGFP and *bmo* (b) is tagged with mCherry. Both Ancestor and *bmo* exhibit uniform biofilm structure. The *fuzY* biofilm is composed of many aggregates. The *wspF* biofilm was in small clusters and was not spatially contiguous on the glass surface, but generated large aggregates in liquid media that were more likely to attach on plastic surfaces (not shown). Mixtures of these mutants were also imaged and are reported in Fig S3.

### Genetic diversity during biofilm selection is maintained by competition and niche differentiation

An important and unusual finding from these studies is that *P. fluorescens* populations grown in the biofilm bead model rapidly and repeatedly diversify into multiple phenotypes and genotypes that coexist, with no genotype ever reaching 100%. This observation implies that selection acts strongly on mutants that inhabit different niches of varied capacity. This leads to the prediction that if putative niche specialists are mixed in equal amounts and propagated, their frequencies should equilibrate according to the capacity of each niche and the relative competitiveness of each mutant in conditions of niche overlap (e.g., growth on a common carbon source). We created a mixed population of the three, representative biofilm-adapted mutants *bmo*, *wspF*, and *fuzY* and their *lac*-marked ancestor. We used this mixed population to inoculate six replicate cultures grown in our bead model of biofilm growth for four days without triclosan and plated daily to determine the frequency of each genotype on newly colonized beads (Fig. 7). As predicted, all genotypes coexisted within each population, with *wspF* and *fuzY* mutants generally at low frequencies and the *bmo* mutant and ancestor at intermediate frequencies. In addition, frequencies of *bmo* and WT were anticorrelated in half of the replicates, which suggests they occupy differentiated but overlapping niches, as the microscopy indicated and ecological literature supports (Fig. 6) (Caplat et al., 2008; Chesson, 2000). We used linear regression to test this hypothesis for all genotype combinations and found a significant, negative relationship for the *bmo* mutant frequency with frequencies of all other genotypes, which is consistent with competition (Table 2, Fig. S4). In summary, our model of the biofilm life cycle exerts strong, parallel selection for different phenotypes that balance biofilm production, growth, dispersal, and reattachment in different ways. Mutants adapted to different aspects of these traits can coexist by invading in conditions in which they are more fit, while declining in conditions where they are less competitive.

**Fig. 7.**
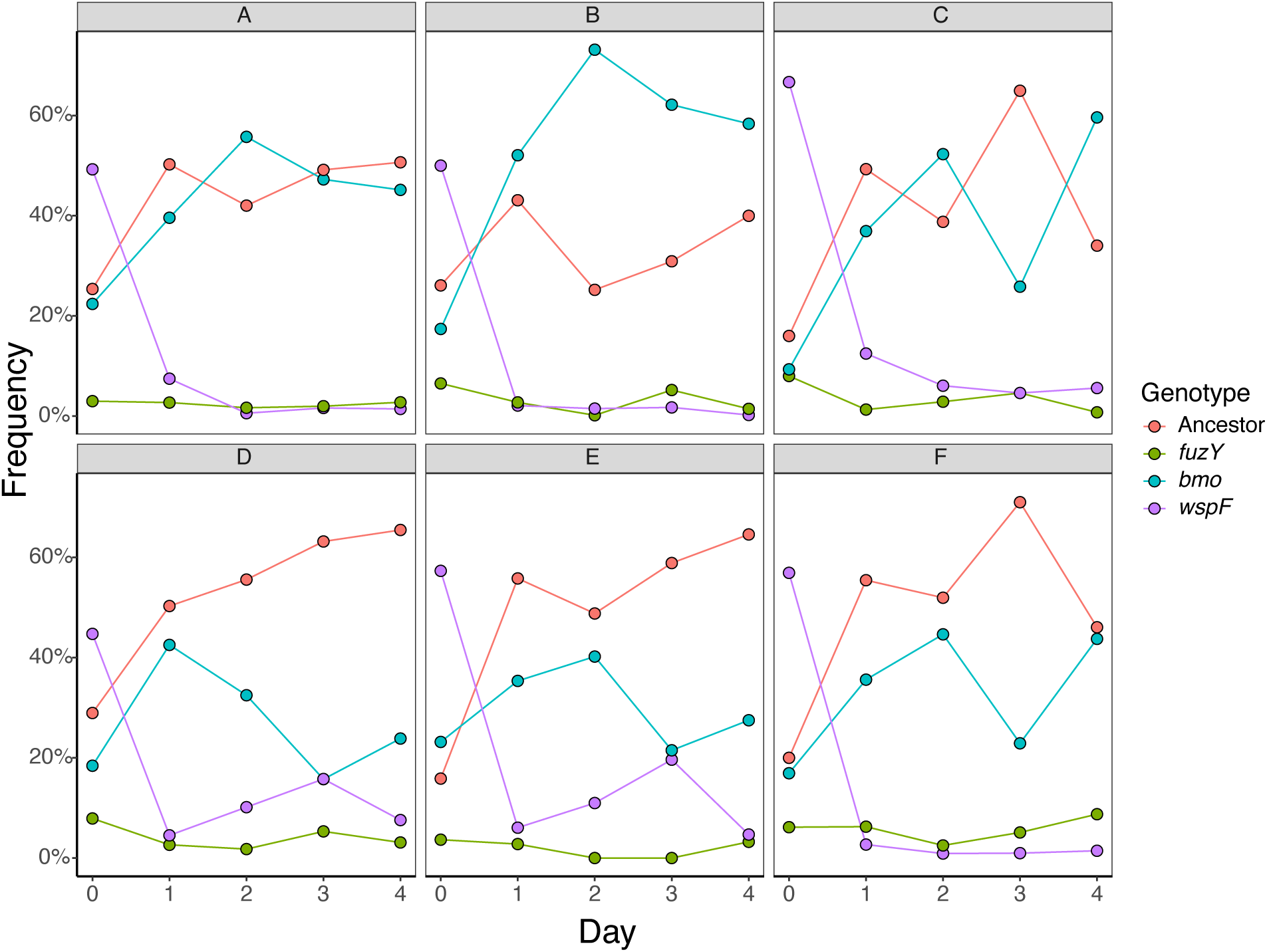
Ecological dynamics of constructed communities indicate multi-niche selection. Six populations containing equal volumes of representative, biofilm-adapted loss-of-function mutants in *bmo* (PFLU0185), *fuzY*, and *wspF* were propagated in the biofilm bead model and plated daily to assess frequencies. All mutants persisted, with *fuzY* and *wspF* declining rapidly in frequency and *bmo* and the ancestor fluctuating at intermediate frequencies and opposite trajectories.

**Table 2.** Linear regressions of frequencies of *bmo* (PFLU0185) between the ancestor (Afreq), *wsp* mutant (Wfreq), and *fzy* mutant (Ffreq), as shown in Figure S4. The significant negative slopes indicate competition.

| Model Equation | Std. Error | t value | R-squared | p-value |
| --- | --- | --- | --- | --- |
| Pfreq = (-0.99)Afreq + 0.91 | 0.07 | -14.69 | 0.79 | <0.0001 |
| Pfreq = (-0.7)Wfreq + 0.46 | 0.32 | -2.22 | 0.08 | <0.05 |
| Pfreq = (-1.84)Ffreq + 0.48 | 0.58 | -3.14 | 0.15 | <0.001 |

## DISCUSSION

### Student experiments discover new genetic pathways of adaptation to the biofilm life cycle

Involving secondary school students in distributed scientific experiments in their own science classrooms offers complementary promises, including: 1) a high level of replication that can broadly test a common hypothesis, and 2) developing a community of teachers and learners who are connected by a common set of research skills, experiences and findings. Here, we not only report evidence that both promises can be met, but also that students contributed to expanding our knowledge of the genetics of adaptation to the biofilm life cycle. Their experiments with the biofilm bead model selected for phenotypes that excel during daily cycles of dispersal, recolonization, and biofilm assembly. By picking many representative clones with these phenotypes and then genotyping them by whole-genome sequencing, we use experimental evolution as a powerful forward genetic screen. This approach identified mutants that support previous findings from pellicle selection experiments and also uncovered novel adaptive pathways within the biofilm lifestyle. These findings support the hypothesis that biofilm selection acts on traits governed by a few central regulators, leading to strong parallel evolution across experiments. They also reveal that different selective environments, for example associated with varied implementation in classrooms, can uncover alternative adaptations in novel pathways to biofilm formation. Because most of these mutants contain only one mutation in their genome, the relationships between genotype and phenotype are direct. In this section, we discuss predicted molecular and biochemical effects of these mutants in detail, several of which strengthen or modify existing models of *Pseudomonas* biofilm gene regulation. We begin by analyzing mutations producing the well-known wrinkly and fuzzy colony morphologies and follow with discussion of the potential causes of beneficial small colony variants.

### Student-selected colony variants support previous findings of genetic pathways important for biofilm formation

The most common cause of wrinkly phenotypes was mutations in the *wsp* pathway, a chemotaxis-like system composed of 7 genes: *wspA-R*, under the control of a single promoter (Fig. 2) (Bantinaki et al., 2007). WspA is a methyl-accepting chemotaxis protein that is anchored at the cytoplasmic membrane by the scaffolding proteins WspB and WspD. Upon sensing an environmental signal, conformational changes in the WspA-B-D complex result in WspA being methylated by WspC, a constitutively active methyltransferase. This begins a signaling cascade that activates WspE, a histidine kinase response regulator. WspE goes on to activate WspR, a DGC, through phosphorylation, increasing levels of cyclic di-GMP and ultimately leading to increased biofilm formation through increased cellulose production (Spiers et al., 2002, 2003). WspE also phosphorylates WspF, a methylesterase that negatively regulates the system by demethylating WspA. In general, the Wsp pathway mutations we identified are predicted to cause constitutive production of cyclic di-GMP and, consequently, increased biofilm formation. We identified deletions in *wspA* that have been shown *in P. aeruginosa* to lock the pathway on in a methylation-independent manner (A. Xu et al., 2021), whereas nonsynonymous mutations in the *wspA* signaling domain likely disrupt the stability of its trimer-of-dimer interactions, increasing WspE activation (O’Connor et al., 2012; Kessler et al., 2021). The mutation producing a WspC-WspD fusion protein likely leads to hypermethylation of WspA because it physically links WspC to the WspA-B-D complex (Kessler & Kim, 2022). The many different mutations in *wspF* include nonsynonymous substitutions, many of which occurred at the same site repeatedly, insertions, and deletions, including frameshift and nonsense mutations that introduced premature stop codons (Fig. 2). These mutations almost certainly interfere with WspF activation and/or its ability to demethylate WspA (O’Neal et al., 2022; A. Xu et al., 2021).

The second most common cause of wrinkly phenotypes were mutations in the *yfiBNR* pathway, which is composed of three genes under the control of a single promoter (Fig. 2). YfiN is a DGC that is localized to the cytoplasmic membrane and negatively regulated by the attachment of the periplasmic YfiR protein (Malone et al., 2010, 2012). In response to membrane stress, YfiB, a lipoprotein located at the outer membrane (Malone et al., 2012), adopts an elongated conformation, which allows it to bind to and sequester YfiR (Li et al., 2018; M. Xu et al., 2016). YfiN is therefore released from its negative regulation and able to catalyze cyclic di-GMP production, promoting biofilm formation (Fig. 2). The C-terminal region of *P. aeruginosa* YfiR is predicted to contain the attachment site to YfiN (Malone et al., 2012; Yang et al., 2015); therefore, the *yfiR* missense and nonsense mutations we discovered in this region may eliminate YfiR’s ability to bind to YfiN. *P. aeruginosa* YfiR forms a dimer in its active form via salt bridges formed by R98 and D80 and hydrogen bonds between T76 (Yang et al., 2015). Notably, five *yfiR* mutants experienced the same 33bp deletion from T75 to T85 in *P. fluorescens* that removes these homologous regions involved in dimerization. In addition, both this deleted region and the downstream 33bp deletion contain or are located close to conserved cysteine residues that in *P. aeruginosa* form disulfide bonds important for proper protein folding (Yang et al., 2015). Deletions in these regions are therefore likely to lead to protein misfolding that inhibits YfiR’s ability to bind YfiN, thereby derepressing the DGC which promotes a biofilm lifestyle.

Mutations in *morA* were the third most likely cause of wrinkly phenotypes. MorA is a single protein localized to the cytoplasmic membrane that contains two PAS sensor domains as well as active DGC and PDE domains, so it can catalyze both production and degradation of cyclic di-GMP (Fig. 2) (McDonald et al., 2009; Phippen et al., 2014). We identified two identical nonsynonymous substitutions in the DGC, three mutations affecting the linker region between DGC and PDE domains that likely cause conformational changes that increase DGC activity (Mhatre et al., 2020), and three identical mutations in the PDE that appear to disrupt its activity (Fig. 2). The elevated biofilm of all these mutants indicates that their effects increase cyclic di-GMP levels.

Fuzzy colony phenotypes are caused by loss-of-function mutations in *fuzY* (Fig. 3). In *P. fluorescens* SBW25, *fuzY* is part of a five-gene operon that was predicted to encode a β-glycosyltransferase that plays a role in modifying lipopolysaccharides (LPS) O-antigens, the outermost portion of the LPS (Ferguson et al., 2013). The reciprocal best match of *fuzY* homologs in *P. aeruginosa* strain PAO1 is *wapH*, which participates in synthesis of the LPS core (Lam et al., 2011). In either case, in addition to roles in pathogenesis and symbiosis, O-antigen structure plays an important role in bacterial adhesion and cohesion (Lerouge & Vanderleyden, 2002). In *P. aeruginosa*, O-antigen mutants exhibit increased cell-cell cohesion and increased adhesion to glass surfaces (Lau et al., 2009). In our bead selection experiment, we expect that mutations in *fuzY* are beneficial because they increase both cell-cell and cell-surface contact, which would provide an advantage in adhering to the surface of the bead.

### Student findings uncover new pathways to biofilm adaptation predicted to disrupt periplasmic protein folding

Biofilm evolution experiments with our bead model often select for small colony variants with increased Congo Red binding, which is consistent with higher cellulose production (Fig. 1; Fig. S5). These phenotypes were consistently caused by mutations in a pathway that regulates the formation of disulfide bonds, *dsbRS* and *dsbDEG*, which is critical for some proteins to be properly folded into their active state conformation (Fig. 3). Most knowledge of enzymes involved in the formation of disulfide bonds comes from studies in *Escherichia coli* (J. C. A. Bardwell et al., 1991; J. C. Bardwell et al., 1993). In *E. coli*, the periplasmic oxidoreductase DsbA catalyzes formation of disulfide bonds but is thought to perform this reaction indiscriminately by introducing bonds between neighboring cysteines (Bader et al., 2000; J. C. A. Bardwell et al., 1991). Proteins that require disulfide bond formation between nonconsecutive cysteines undergo further processing through the activity of disulfide isomerases (DsbC and DsbG) that rearrange these bonds (Berkmen et al., 2005; Hiniker & Bardwell, 2004; Manta et al., 2019).

In *P. aeruginosa*, homologs of the regulatory two-component system *dsbRS* have been shown to regulate the activation of *dsbDEG* homologs partly by binding copper ions (Yu et al., 2022). Based on this study, we predict that evolved *dsbS* mutants lead to altered phosphorylation of *dsbR* which interferes with its regulation of *dsbDEG*. The evolved intergenic mutation is found in the predicted DsbR binding site and would therefore reduce *dsbDEG* transcription. The evolved DsbE-DsbG fusion mutant likely interferes with the ability of these proteins to fold properly, with the downstream consequence of misfolded periplasmic proteins. Taken together, we predict that all mutations in this pathway result in misfolded periplasmic proteins, although the exact proteins that are affected are unknown. Interestingly, in *P. aeruginosa*, deletion of *dsbA* led to a small colony variant phenotype that was hypothesized to be a result of YfiR misfolding due to its cysteine residues that contribute to dimerization, resulting in constitutive activation of YfiN and elevated cyclic-di-GMP (Malone et al., 2012). However, the small colonies produced by *dsb* mutations differ from the wrinkly mutants of caused by *yfiR* mutations, suggesting different effects on biofilm requiring further study.

### Discovery of frequent mutations in phosphodiesterase *bmo* that do not change colony morphology

Clones retaining the ancestral, smooth colony phenotype at the end of evolution experiments often acquired a mutation in *bmo*, a gene with both DGC and PDE domains that is highly conserved in the genus *Pseudomonas* and predicted to function primarily as a PDE (Fig. 3) (Eilers, Hoong Yam, et al., 2024; Wei et al., 2019). A representative mutant showed intermediate levels of both biofilm production and motility, leading us to name this gene *bmo* for its role in balancing both biofilm and motility. We predict that our model selects for *bmo* mutants because they act as biofilm generalists that have improved attachment and biofilm formation compared to their ancestor, while also maintaining the ability to disperse from the old bead to recolonize the new bead. Characterization of multiple biofilm-associated phenotypes supports this hypothesis. Compared to the ancestor and the hyper-biofilm *wspF* mutant, *bmo* has intermediate motility, cyclic di-GMP, and overall biofilm attachment phenotypes, whereas its cellular attachment to the bead is as high as *wspF* (Fig. 5). Moreover, *bmo* biofilm phenotypes are nearly identical to the LPS mutant *fuzY*, although *bmo* has somewhat improved motility, attachment, and overall biofilm formation, along with significantly greater cyclic di-GMP levels. It is also notable that *bmo* mutants appear to be selected more frequently in media containing triclosan, suggesting the stress imposed by this compound interacts with the *bmo* regulatory pathway that remains to be defined.

*P. aeruginosa bmo* homologs have been shown to influence biofilm formation and regulate basal levels of cyclic di-GMP. A deletion mutant of the *P. aeruginosa* PA14 homolog (PA14_03720) had reduced swimming motility and increased twitching motility in comparison to wild-type PA14 (Ha et al., 2014). Deletion mutants of the *P. aeruginosa* PAO1 homolog (PA0285, *pipA*) have increased biofilm formation, aggregation, Psl production, and cyclic di-GMP production, and decreased swimming motility (Cai et al., 2022; Eilers et al., 2022; Wei et al., 2019). A recent study that compared deletion mutants of *bmo* homologs in *P. aeruginosa* strains PA01, PAK, and PA14, and *P. putida* strain KT2440 found that these mutants had increased early attachment (at 4, 6, and 8 hours) in comparison to their wild-type counterparts and that the magnitude of these differences depended on strain background (Eilers, Hoong Yam, et al., 2024). Among the 22 unique mutants detected by deep sequencing populations evolved in the bead biofilm model, most are predicted to (1) cause loss-of-function by introducing premature stop codons or (2) negatively affect PDE function or PAS domain signaling through nonsynonymous SNPs (Sup. Table 1). We predict these mutants resemble the representative mutant clone we studied (Fig. 5) by having a higher basal level of cyclic di-GMP that improves their attachment to the bead surface and biofilm formation.

### Phenotypic differentiation among mutants reflects niche differentiation that enables coexistence

The spectrum of biofilm phenotypes arising in these experiments suggests that they exploit different ecological conditions, which could explain both their rapid emergence and subsequent coexistence. This model was well supported by confocal microscopy of biofilms produced by individual mutants and mixed pairs (Fig. 6, Fig. S3) but these measurements were on glass rather than polystyrene and did not allow mutants to reach equilibrium frequencies. Therefore, we mixed representative mutants of each morphology and tracked their frequencies over a week of passage. The *bmo* mutant and the ancestor appeared to compete as their intermediate frequencies oscillated, whereas *wspF* and *fuzY* mutants rapidly declined but were maintained at low (∼5-10%) frequencies (Fig. 7). The cycling between the *bmo* mutant and the ancestor can be explained by their occupying overlapping but distinct niches (Brisson, 2018; Chesson, 2000), whereas the persistence of *wspF* and *fuzY* at lower frequencies suggests they inhabit more differentiated biofilm niches but are outcompeted by the other genotypes under most conditions (Fig. 6, Fig. S3, Table 2). These dynamics agreed with the phenotypic dynamics from the 15-day evolution experiment (Figure 4), in which biofilm-adapted mutants with distinct wrinkly and fuzzy colony morphologies rapidly arose and persisted at low frequencies while ancestral colony morphologies consistent with persistence of the ancestor and biofilm-adapted *bmo* mutants predominated. These overall interactions conform to a model known as multi-niche selection (Levins, 1963), which requires that each species has a selective advantage in some habitats while others are superior in other habitats, with areas of niche overlap. We suggest that the repeated emergence and maintenance of genotypes with differentiated niches within the biofilm life cycle provides a clear example of how variable environments can favor balanced polymorphisms (Martin et al., 2016; Poltak & Cooper, 2011). Given the ubiquity of microbial biofilms undergoing regular cycles of attachment, aggregation, dispersal, and reattachment, we can expect genetic and phenotypic heterogeneity like that shown here to be widespread. This diversity may therefore be expected in biofilms produced by natural populations of *P. fluorescens* in and around plants or in those produced by its relative *P. aeruginosa* in the built environment or when establishing opportunistic infections.

### Building a community of learners studying biofilm evolution who understand its biomedical relevance

We began the EvolvingSTEM project in 2013 with the aim of teaching evolution and genetics in high school classrooms through an authentic bacterial evolution experiment. Our rationale was that the predominant way of teaching natural selection through passive, historical examples of speciation would be less effective than the experience of seeing evolution in action in student-led experiments. We showed that high school students achieved greater learning of content meeting Next Generation Science Standards (States, 2013) in a prior study (Cooper et al., 2019). But this work also held promise for expanding our understanding of mechanisms of biofilm adaptation, given the increasing speed and economy of genotyping mutants by whole-genome sequencing. As we collected these mutants and identified their genetic causes, we added these discoveries into our curriculum materials and emphasized how these mutants reinforced and extended our understanding of how *P. fluorescens* and its pathogenic relative *P. aeruginosa* adapt when biofilm formation is advantageous. Mutations in the *wsp* cluster, in *yfiBNR*, and in *morA* are commonly recovered in infections caused by *P. aeruginosa* (Gloag et al., 2019; Malone et al., 2010; Marvig et al., 2015; Smith et al., 2006), and their recovery here indicates their centrality to this lifestyle switch. In addition, our discovery of mutants in *fuzY* adds to evidence that LPS alterations enhance aggregation and potentially alter susceptibility to bacteriophage (Ferguson et al., 2013), and mutants in the *dsb* pathway and *bmo* highlight less understood genetic pathways to biofilm production that warrant more research into how *Pseudomonas* senses surfaces and initiates attachment.

The experience of offering this curriculum in subsequent years at more than 30 middle and high schools suggests broader and potentially more potent benefits, including improved student attitudes towards science and technology and greater student confidence as science learners, which we have and will report elsewhere (Quigley et al., 2025). One reason that students may begin to see themselves as scientists is the peer reinforcement from collaborating in small groups to conduct the experiments and as an entire class to discuss their findings. Likewise, often multiple classes per school participate in EvolvingSTEM, and within Pittsburgh Public Schools, twelve different schools conducted these experiments in 2024. This article synthesizes their discoveries and we hope encourages greater sharing of student discoveries among more than 2500 students annually who learn about the medical and natural relevance of microbial biofilms from our curriculum. They complete their experiments knowing that the mutants they identify may contribute to a greater understanding of how bacteria evolve in biofilms and ultimately how to manage these evolving communities.

## MATERIALS & METHODS

### School bacterial growth conditions

Students cultured *Pseudomonas fluorescens* strain SBW25, a plant-colonizing bacteria that was isolated in 1989 from the leaf surface of a sugar beet plant grown at the University Farm (Whytham, Oxford, UK) (Bailey & Thompson 1992; Rainey & Bailey 1996 Molecular Microbio). Students cultured bacteria in test tubes that contained 5mL of Queen’s B growth media (Cooper et al. 2019, 20g Proteose Peptone No. 3, 1.5g Potassium Phosphate Dibasic, 25mL Glycerol, and 6mL 1M Magnesium Sulfate per liter) and a polystyrene bead and plated their bead-attached bacterial populations on Half-Strength Tsoy-Agar plates (15g Tsoy and 15g Agar per liter) at the beginning and end of the experiment. In addition, 25μg/mL of the broad-spectrum antimicrobial triclosan was added to the media and agar plates used in the experiments at SciTech and Allderdice to inhibit bacterial and fungal contamination. *Pseudomonas* species are naturally resistant to triclosan, and this concentration was chosen because it is used in *Pseudomonas* isolation agar (e.g., Cat# 17208 Pseudomonas Isolation Agar from Millipore Sigma).

WHS students grew *P. fluorescens* cultures for 13 days at 28 **°**C on a roller drum under biofilm selection conditions, which included 7 bead transfers. SciTech and Allderdice students grew cultures for 4 days on an orbital shaker at 150rpm under biofilm selection conditions, which included 3 bead transfers. SciTech students grew their cultures at 28 **°**C, while Allderdice students grew their cultures at room temperature because they did not have an available incubator.

### Genome sequencing and identification of mutants

Clones were selected from the final plating day of the EvolvingSTEM classroom experiment and grown overnight in 5mL QB media. DNA was extracted from 1mL of the overnight culture with the DNeasy Blood and Tissue Kit (Qiagen). Illumina whole-genome sequencing was performed by our laboratory to a minimum depth of 40x coverage on an Illumina NextSeq500 using the methods described in Baym et al. 2015. Trimmomatic (Bolger et al. 2014) was used to remove sequence adaptors, along with low-quality sequence fragments. We identified mutations by comparing evolved clones to the published genome of *P. fluorescens* SBW25 (NCBI Reference Sequence: NC_012660) using breseq (Deatherage and Barrick 2014). All sequencing data are available at NCBI Bioproject PRJNA1284392.

The ancestral *P. fluorescens* SBW25 populations all exhibited a smooth colony morphology phenotype. In comparison to the published genome sequence, the ancestor used by all student populations had three intergenic mutations present in their ancestral populations: (1) +G at position 45,881; (2) +C at position 985,333; and (3) C_5→3_ at position 3,447,984. The ancestor used by WHS in 2015 also had a 129bp deletion at position 360,368 and an 132bp deletion at position 6,223,020.

### Experimental evolution and longitudinal whole-population sequencing

Populations were initiated with a single *P. fluorescens* SBW25 ancestral clone and evolved for 15 days (14 transfers) in 2mL Queen’s B growth medium at 28**°**C on an orbital shaker at 150rpm. Four replicate lineages were propagated under two biofilm conditions: biofilm selection + 25μg/mL triclosan and biofilm selection without triclosan. Biofilm lineages were propagated using four beads each (following the methods of (Santos-Lopez et al., 2019; Scribner et al., 2020). After 24 hours growth, one bead was transferred to a new test tube with three new beads.

To perform whole-population sequencing, we collected bead-attached bacteria on days 3 and 15 by transferring two beads from each population to a tube with 1.2mL PBS, which was vortexed for 2 minutes to release bacteria from the beads. 500uL was immediately centrifuged for 10 minutes at 14,000rpm, the supernatant was removed, and pellets were stored at −20C for DNA extraction. Sequencing libraries were prepared to a minimum read depth of 150x coverage and analyzed as above with the additional “-p” polymorphism flag in breseq to characterize all reliable mutations at intermediate frequencies in these populations.

### Motility Assays

Assays were performed as described in Filloux and Ramos (2014). Images were analyzed with FIJI (Schindelin et al., 2012). For swarming motility, we grew 6-hour cultures in LB-Lennox Broth (10g Tryptone, 5g Yeast Extract, 5g NaCl per L) at 28℃, 150rpm, then inoculated Swarming Motility Agar Plates (0.6% agar, 1x M8 solution, 0.2% glucose, 0.5% casamino acids, 1mM MgSO_4_) with 2.5μL culture pipetted onto the center of the plate, at the surface of the agar. Plates were incubated upright at room temperature (∼22℃) for 16-24 hours and photographed. For swimming motility, we grew 6-hour cultures of *P. fluorescens* in LB-Lennox Broth at 28C, 150rpm, transferred 100μL to a 1.5mL microcentrifuge tube, dipped a 200uL pipette tip into the culture and used this to stab into the center of Swimming Motility Agar Plates (same recipe as Swarming Motility Plates, but with 0.3% agar). Plates were incubated upright at room temperature (∼22℃) for 16-24 hours and photographed. For twitching motility, we stabbed a match-head sized inoculum from a 2-day old streak plate into the center of an LB-Lennox Agar Plate (10g Tryptone, 5g Yeast Extract, 5g NaCl, 10g Agar, 0.5g Tetrazolium Red, per L) all the way down to the plastic. Plates were inverted and stored in stacks of 3 or less in a plastic box lined with wet paper towels at 28℃ with box lid slightly ajar for 24 hours and photographed.

### Biofilm assays

Our biofilm assay is based on the bead model developed by our lab for studying biofilm formation and evolution. Bacterial cultures were inoculated in triplicate in 2 mL of Queen’s B media and two plastic beads (Cospheric, Polystyrene Spheres, 7.00mm +/- 0.025mm) and grown overnight at 150 rpm on an orbital shaker at 28℃. Each culture was sonicated for 10 seconds at 30% amplitude and used to inoculate two new cultures at a 1:100 dilution in QB both with and without 30 μg/mL triclosan and two polystyrene beads. Cultures were then grown for 24 hours on an orbital shaker at 28℃, 150 rpm, serial diluted, and plated. Biofilm formation was quantified by transferring one of the two beads to a single well of a 24-well plate with 1.5mL of an 0.1% crystal violet solution and incubated at room temperature for 15 minutes. The bead was then washed three times in water for one minute each time to remove excess crystal violet.

Biofilm-bound crystal violet was then solubilized by vortexing the bead for 2 minutes in a 2mL tube filled with 1mL solubilization solution: 95% ethanol, 4.95% water, 0.05% Triton X-100. The concentration of crystal violet was measured at an absorbance wavelength of 590 nm (OD590). Total colony forming units per bead were determined by serial diluting and plating bacteria attached to the second bead. We adopted this method to (1) more accurately represent the conditions in which we grow biofilms in our experiments and (2) allow us to differentiate between total biofilm production and bacterial attachment to the bead. As an indicator of EPS production, we plated mutants on half-strength T-Soy Agar plates (15g Tsoy, 15g Agar, 0.02g Congo Red, 0.0075g Brilliant Blue per liter) and visually assessed dye uptake, where dark red colonies indicate EPS overproduction.

### Intracellular cyclic di-GMP Assay

The *P. fluorescens* ancestor and biofilm-adapted mutants were transformed with the cyclic di-GMP reporter plasmid, pCdrA-gfpC (Addgene plasmid #111614) (Rybtke et al., 2012). The reporter transcriptionally fuses the *P. aeruginosa* PA01 *cdrA* promoter, which is responsive to cyclic di-GMP, to genes encoding green fluorescent protein. Starter cultures of each transformed *P. fluorescens* genotype and an untransformed ancestor control were grown in their own cultures of 2 mL of Queen’s B media at 28℃ on an orbital shaker at 150 rpm for 24 hours. Following this, experimental cultures containing 4 mL of Queen’s B media were inoculated with 10 μL of a given starter culture and incubated at 28℃ on an orbital shaker at 180 rpm for 24 hours. After incubation, a 100 μL sample of each experimental culture was used to capture fluorescence intensity and OD_600_. Cyclic di-GMP levels at 24 hours were calculated by ln(RFI)/ln(ROD_600_), where RFI is relative fluorescence intensity (the ratio of absolute fluorescence intensity of experimental to control culture), and ROD_600_ is relative optical density (the ratio of absolute optical density at a wavelength of 600 nm of a bacterial culture to fresh media).

### Construction of fluorescently tagged *P. fluorescens* strains

A single colony of *P. fluorescens* was inoculated into 5 ml QB and grown overnight to stationary phase. One mL of culture was pelleted by centrifugation and washed twice with 300 mM sucrose. After resuspending the pellet in 100 uL of 300 mM sucrose, 5 uL pTNS3 and 5 uL of the desired pBT plasmid (Table S3) were added and the suspension was incubated at room temperature for 5 minutes. The suspension was then transferred to an electroporation cuvette (2 mm gap) and pulsed at 2.5 kV. Cells were recovered in 1 mL LB and transformants were selected on LB agar + 30 ug/mL gentamicin at 28°C.

### Confocal microscopy

*P. fluorescens* strains expressing eGFP or mCherry were pre-cultured for 24 hours in glass-bottom plates containing 2 mL of Queen’s B media and shaken on an orbital shaker at a speed of 125 rpm. Biofilms generated by *P. fluorescens* were expected to attach to the surface of the glass. After pre-culture, the old media in the plates were removed by pipetting, and the biofilms were washed twice with 1 mL of PBS to remove excess planktonic bacterial cells. Following the washing step, 1 mL of fresh PBS was added to each glass-bottom plate to maintain the survival and limit the growth of bacterial cells within the biofilm. Images were captured at 400× magnification in both eGFP and mCherry channels on an Olympus FV-1000 confocal microscope. Z-stack images spanned from the bottom to the top of the biofilms. Image processing was conducted using ImageJ (Schindelin et al., 2012) and BiofilmQ (Hartmann et al., 2021).

### Multi-genotype competition assay

We created mixed populations of bacteria by combining a representative mutant of *bmo* (PFLU0185) (with a single nucleotide insertion resulting in a premature stop codon truncating both the DGC and PDE domains), wrinkly (*wspF* Q185*) and fuzzy (*fuzY* Q11*) mutants, along with a lac-marked *P. fluorescens* SBW25 ancestor. All four clones were used to inoculate cultures containing 2mL QB and 2 beads and grown overnight at 28**°**C on an orbital shaker at 150rpm. Both beads were then transferred to 1.5mL QB in small glass tubes and vortexed for 2 minutes and this mixture was used to make freezer stocks. Bacteria from a portion of one freezer stock was diluted in QB and 50uL of this dilution was transferred to 6 replicate populations containing 2mL QB and 4 beads. Starting frequencies of each mutant and ancestor were determined by diluting a spread-plating from each starting culture. Populations were then evolved for 4 days (3 transfers) in 2mL Queen’s B growth medium at 28**°**C on an orbital shaker at 150rpm. Every 24 hours, one bead was transferred to a new test tube with three new beads and all newly colonized beads were vortexed for 2 minutes, diluted, and spread-plated to count changes in colony phenotypes over time.

## ACKNOWLEDGEMENTS

We thank Tyler McAloon and Erin Nawrocki for technical support. We appreciate the commitment of dozens of teachers and thousands of students to implement this curriculum, isolate these mutants, and inspire this research. This work was supported by SEPA R25AI180989 from the NIH, by BIORETS DBI-2147075 from the NSF, and by the Dean’s office from the University of Pittsburgh, School of Medicine.

---

**Table S1.** Complete details of mutant genotypes, including secondary mutations (XLS).

**Table S2.** Details of mutations detected during the evolution experiment using population-wide whole-genome sequencing (XLS)

**Table S3.**
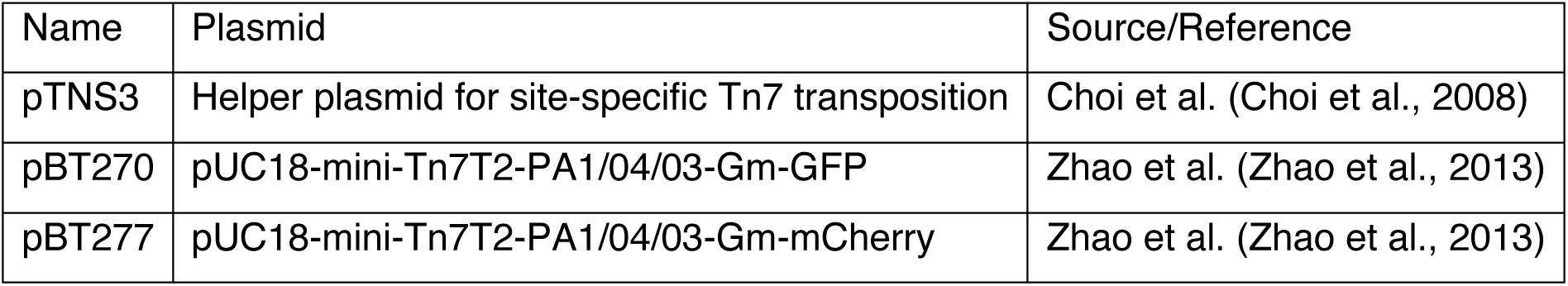
Plasmids used in this study.

| Name | Plasmid | Source/Reference |
| --- | --- | --- |
| pTNS3 | Helper plasmid for site-specific Tn7 transposition | Choi et al. (Choi et al., 2008) |
| pBT270 | pUC18-mini-Tn7T2-PA1/04/03-Gm-GFP | Zhao et al. (Zhao et al., 2013) |
| pBT277 | pUC18-mini-Tn7T2-PA1/04/03-Gm-mCherry | Zhao et al. (Zhao et al., 2013) |

## Figure Legends

**Fig. S1.**
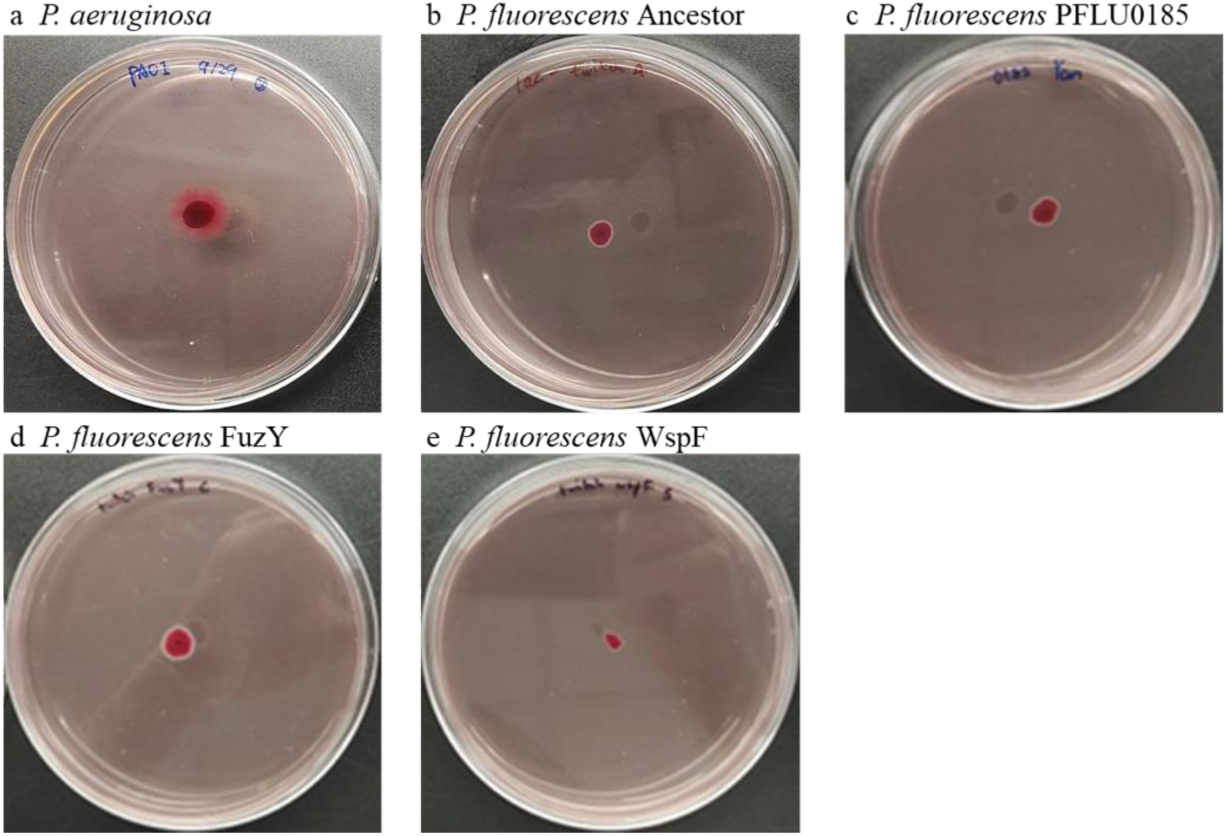
*Pseudomonas fluorescens* lacks twitching motility. Twitching motility was measured on LB-Lennox Agar Plates after 24 hours of growth. PA01 (A) exhibits a twitching phenotype, indicated by the halo surrounding the dark red colony in the center of the plate where inoculation occurred. *P. fluorescens* ancestor (B), *pflu0185* mutants (C), *fuzY* mutants (D), and *wspF* mutants (E) show no signs of twitching motility as all four genotypes lack a halo surrounding the inoculation point.

**Fig. S2.**
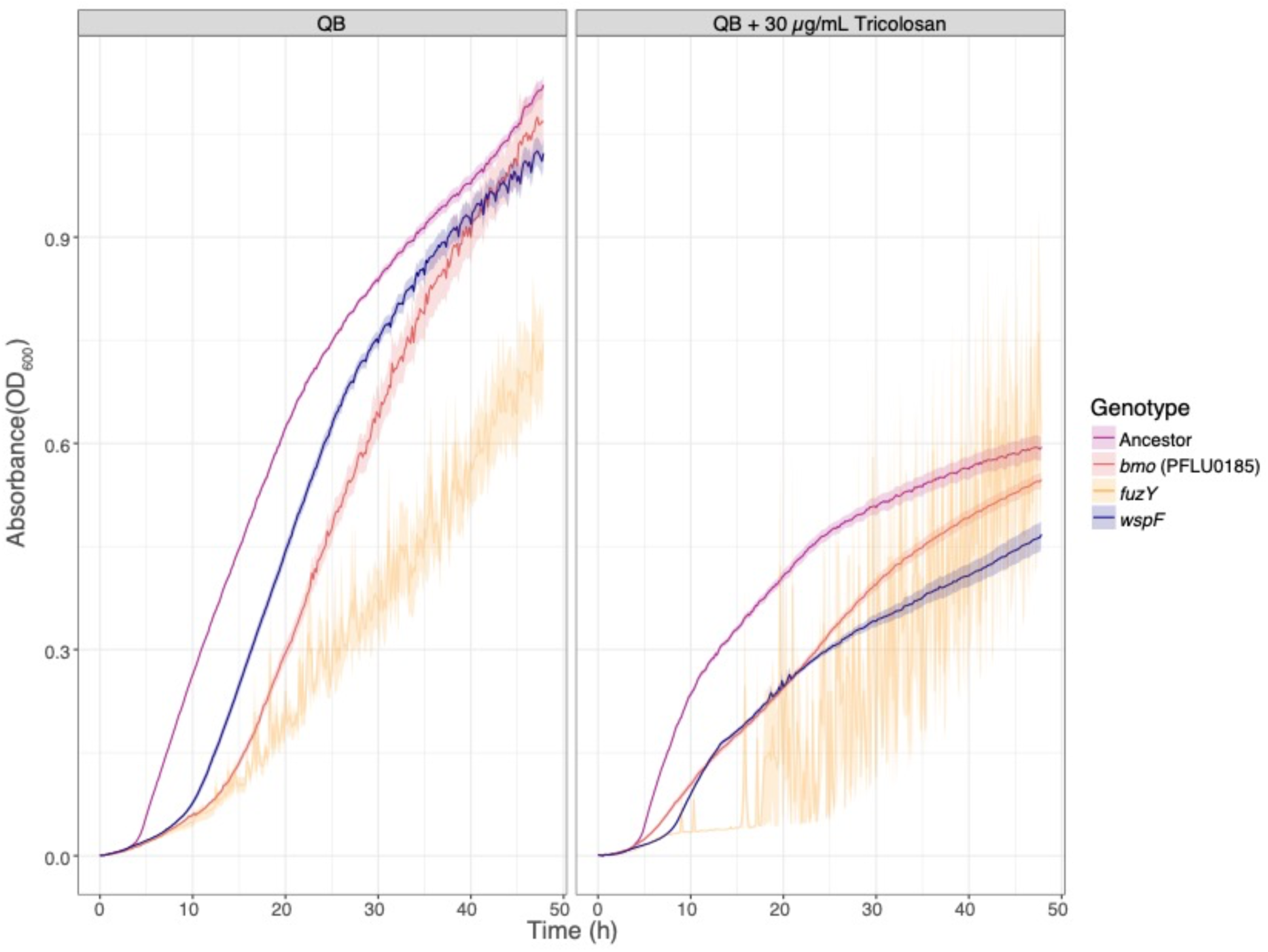
Planktonic growth is inhibited by triclosan.

**Fig. S3.**
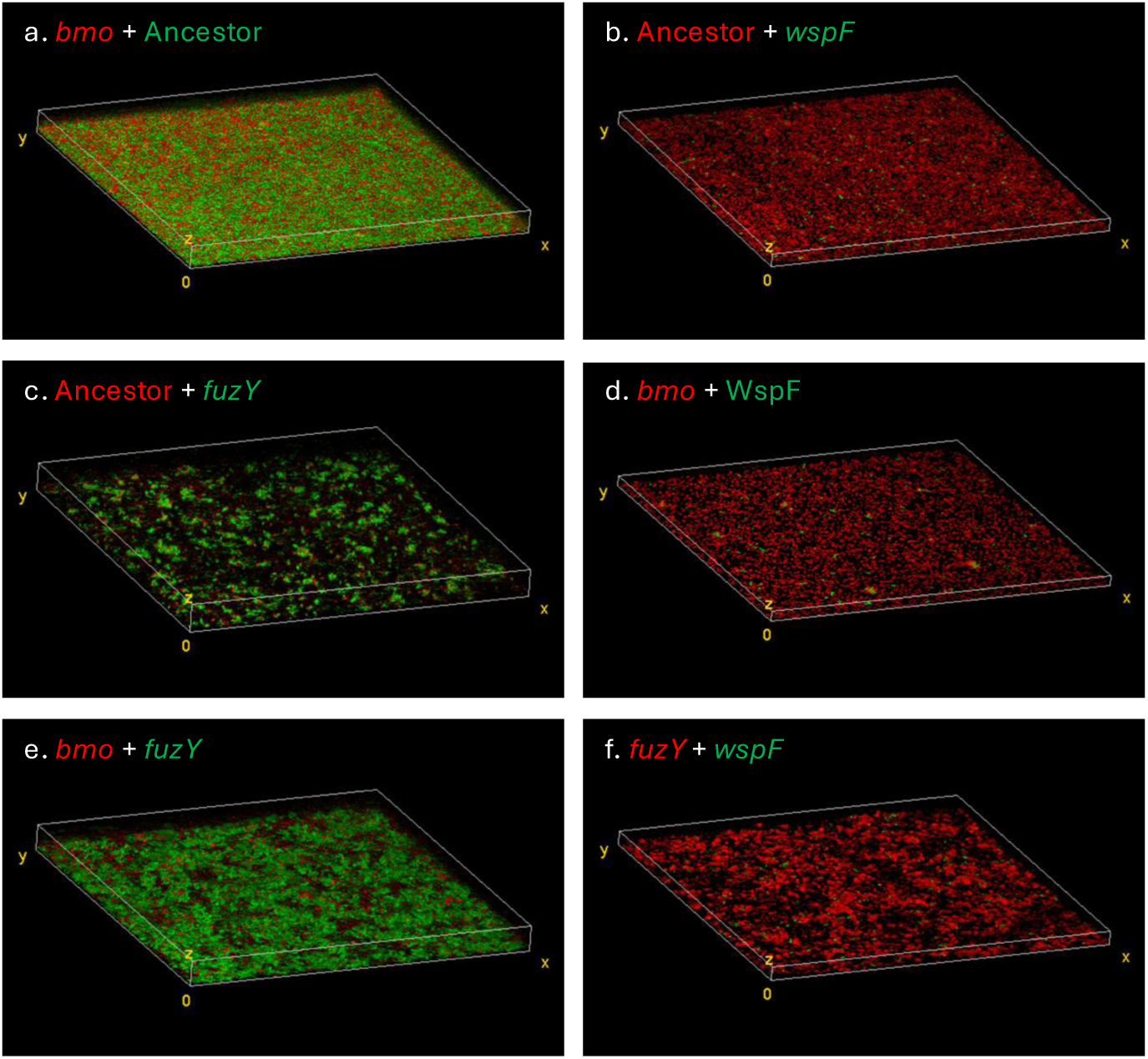
Confocal microscopy of representative *P. fluorescens* mutants mixed in pairs shows differentiated biofilm phenotypes. Methods replicate those of Fig 6. Text color of labels indicates fluorescent marker in that genotype; red = mCherry, green = eGFP.

**Fig. S4.**
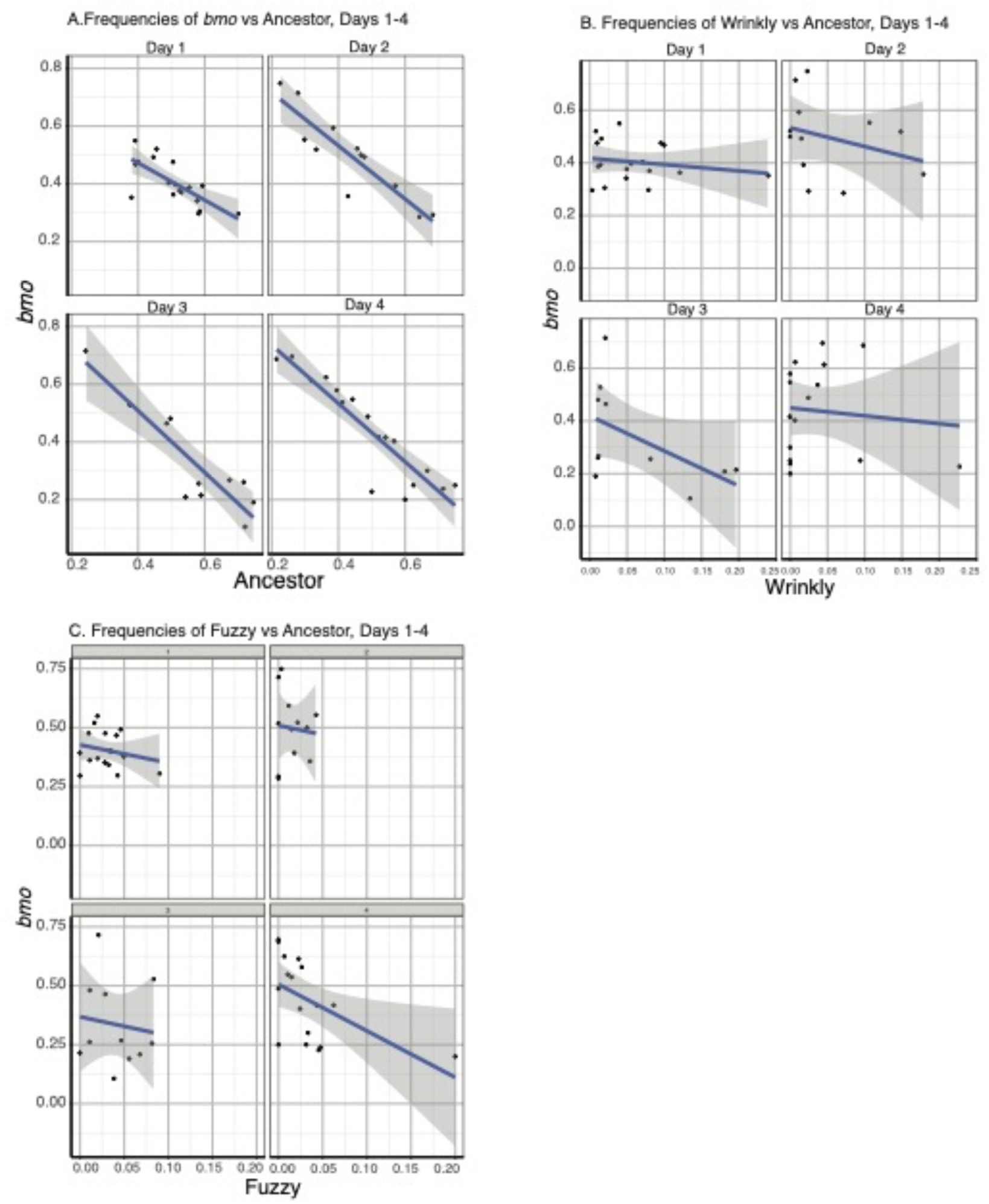
Linear regressions of *bmo* frequencies, as compared to ancestor (A), *wspF* (B), and *fuzY* (C).

**Fig. S5.**
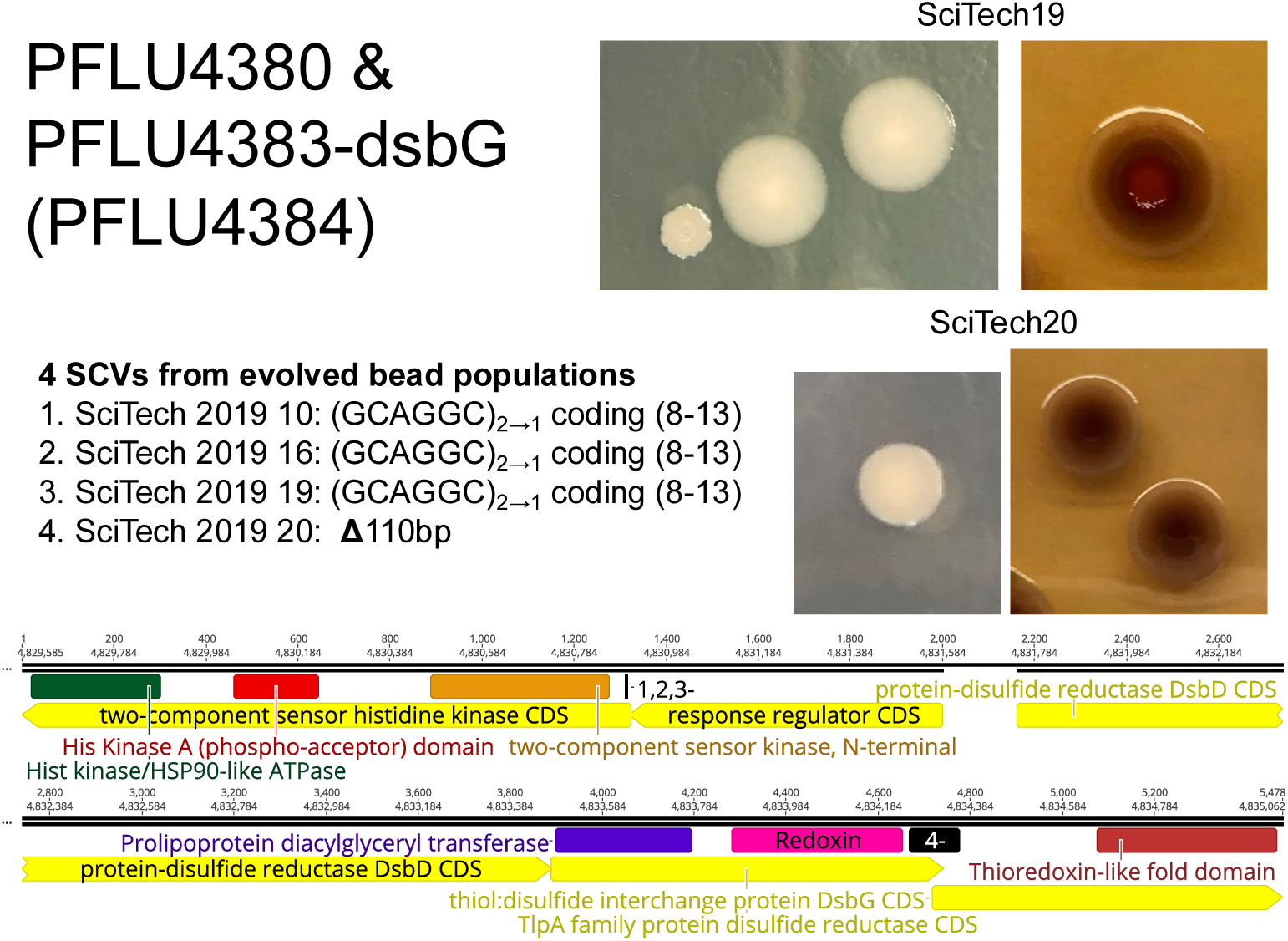
Colony morphology of small colony variants with mutations in *dsb* genes and their differential uptake of Congo Red dye.

## Notes

### Competing Interest Statement

The authors have declared no competing interest.

### Summary of Updates

revised text, updated Figures, clarified methods

